# Metabotyping of Andean pseudocereals and characterization of emerging mycotoxins

**DOI:** 10.1101/2022.06.23.497323

**Authors:** Pedro G. Vásquez-Ocmín, Guillaume Marti, Alice Gadea, Guillaume Cabanac, Juan A. Vásquez-Briones, Sandro Casavilca-Zambrano, Nadia Ponts, Patricia Jargeat, Mohamed Haddad, Stéphane Bertani

**Affiliations:** UMR 152 PHARMADEV, IRD, UPS, Université de Toulouse, Toulouse, France; International Joint Laboratory of Molecular Anthropological Oncology, IRD, INEN, Lima, France; Laboratoire de Recherche en Sciences Végétales (UMR 5546), CNRS, Université de Toulouse, Toulouse, France; MetaboHUB, National Infrastructure of Metabolomics and Fluxomics, Toulouse, France; UMR 5505 IRIT, CNRS, INP, UPS, Université de Toulouse, Toulouse 31400, France; Facultad de Ingeniería Agrícola, Universidad Nacional Agraria La Molina, Lima, Peru; Faculdad de Ciencias de la Salud, Universidad de Huánuco, Huánuco, Peru; Banco de Tejidos Tumorales, Instituto Nacional de Enfermedades Neoplásicas, Lima, Peru; UR 1264 MYCSA, INRAE, Villenave d’Ornon, France; UMR 5174 EDB, CNRS, IRD, UPS, Université de Toulouse, 31062 Toulouse, France

**Keywords:** Andean pseudecereals, metabotyping, mycotoxins, fungal communities

## Abstract

Pseudocereals are best known for three crops derived from the Andes: quinoa (*Chenopodium quinoa*, Chenopods I), canihua (*C. pallidicaule*, Chenopods I), and kiwicha (*Amaranthus caudatus*). Their grains are recognized for their nutritional benefits; however, there is a higher level of polyphenism and the chemical foundation that would rely with such polyphenism has not been thoroughly investigated. Meanwhile, the chemical food safety of pseudocereals remains poorly documented. Here we applied untargeted and targeted metabolomics approach by LC-MS to achieve both: **i**) a comprehensive chemical mapping of pseudocereal samples collected in the Andes to classify them according to their chemotype; **ii**) a quantification of their contents in emerging mycotoxins. An inventory of the fungal community was also realized with the aims to better know the filamentous fungi present in these grains and try to parallel this information with the presence of the molecules produced, especially mycotoxins. Metabotyping permitted to add new insights into the chemotaxonomy of pseudocereals, confirming the previously established phylotranscriptomic clades: Chenopods I (clusters quinoa and canihua), and Amaranthaceae s.s. (cluster kiwicha). Moreover, we report for the first time the presence of mycotoxins in pseudocereals. Sixteen samples of Peru (out of 27) and one sample from France (out of one) were contaminated with Beauvericin, an emerging mycotoxin. There were several mycotoxigenic fungi detected, including *Aspergillus sp*., *Penicillium sp*., and *Alternaria sp*., but not *Fusaria*.

**Highlights:**

▪ Twenty-seven grain samples of Andean pseudocereals were profiled by LC-HRMS.
▪ Untargeted metabolomics was used to differentiate varieties from the whole metabolome dataset.
▪ Five mycotoxins were quantify using targeted metabolomics.
▪ Sixteen samples of Peru and one sample from France were contaminated with Beauvericin.
▪ An inventory of the fungal community infesting the Andean pseudocereal samples was documented.

## 1. Introduction

Pseudocereals are best known for three crops derived from the Andes: quinoa (*Chenopodium quinoa*), canihua (*C. pallidicaule*), and kiwicha (*Amaranthus caudatus*). The use of these pseudocereals as “superfood” for healthier diets has been growing exponentially over the past decade because of their high content of essential amino acids, antioxidants, fibers, and vitamins while being gluten-free (Pereira et al., 2019). Due to its high nutritional value, the Food and Agriculture Organization (FAO) has deemed quinoa a “golden seed” because of its potential to ensure food safety and nutrition for the world’s population amid climate change. Thus, pseudocereals, are not only recognized for their nutritional benefits but also for their ability to adapt to a variety of agro-environmental conditions. Indeed, Peru, Bolivia, and Ecuador are the main producers of quinoa, canihua, and kiwicha, where these pseudocereals are especially well adapted to the abiotic stresses found in Andean highlands (Bonifacio, 2019).

There is a higher level of polyphenism among the pseudocereals cultivated in the Andean countries. For instance, three types of quinoa have been categorized according to the color of the seeds: red quinoa, white quinoa, and black quinoa. However, the chemical foundation that would rely upon such polyphenism has not been thoroughly investigated. Meanwhile, the chemical food safety of pseudocereals remains poorly documented as well. Despite their appeal to an organic diet, pseudocereals may still have the potential for adverse effects from secondary metabolites produced by infesting microorganisms, such as mycotoxin-producing fungi. Mycotoxins are naturally occurring secondary metabolites with adverse effects on human and animal health upon ingestion. They are mainly produced by strains of the three fungal genera *Aspergillus, Fusarium*, and *Penicillium* (Bräse et al., 2009). More than 25% of the crop production worldwide bears contamination with mycotoxins, making them ubiquitous in cereals and cereal-derived foods (Eskola et al., 2020; Lee & Ryu, 2017). Therefore, it is essential to better understand local diversity and mycotoxigenic risk for pseudocereals in the context of a global demand that encourages market diversification and sustainable and equitable development.

Metabolomics based on technology like hyphenated liquid chromatography-mass spectrometry (LC-MS) assisted by bioinformatics and statistical analysis is a powerful approach for both the detection and the annotation of key compounds, as well as for deciphering their biological functions. Our team has previously implemented a workflow based on untargeted LC-MS-based metabolomics able to achieve the metabotyping of plants by identifying their main biomarkers (Vásquez-Ocmín et al., 2021). In the present study, we applied this approach to both achieve a comprehensive chemical mapping of pseudocereal samples collected in the Andes, and to classify them according to their chemotype. Additionally, targeted LC-MS metabolomics was applied on the same samples to quantify their contents in mycotoxins Aflatoxin B_1_, Beauvericin, Fumonisin B_1_, Ochratoxin A, and Patulin. Finally, an inventory of the fungal community infesting pseudocereal samples from the Andes was documented, thereby correlating the presence of toxicogenic fungi and their associated secondary metabolites.

## 2. Material and methods

All solvents used for chromatography were LC-MS grade (HiPerSolv Chromanorm, VWR, Fontenay-Sous-Bois, France). Milli-Q RG system (Millipore, France) was used to produce high purity water of 18.2 MΩ.cm resistivity. Formic acid was >98% for LC-MS (Fluka, Buchs, Switzerland). Mycotoxins standards [*i*.*e*., Aflatoxin B_1_ (AflaB_1_), Beauvericin (BEA), Fumonisin B_1_ (FumB_1_), Ochratoxin A (OTA), and Patulin (PAT)] were purchased from Sigma-Aldrich (St. Louis, MO, USA) and immediately stored at -30 °C.

### 2.1. Plant material

Twenty-seven grain samples were collected between September 2019 and December 2020 in seven regions of Peru (elevation range: 1,410-3,855 meters height above mean sea level) and one grain sample was bought in a local market in Toulouse, France. For quinoa samples from Peru, eight grains were collected directly (unwashed) and the 13 others having been washed (see the list and description of varieties in **supp info 1**). After the collection, all grain samples were immediately stored in airtight bags at -20°C. Authorization to access genetic resources was granted by the competent authority (INIA) under Biodiversity Collection Permit Number 021-2018-MINAGRI-INIA-DGIA/SDRIA.

### 2.2. Metabolome analyses

#### 2.2.1. Untargeted metabolomics

For each grain sample (100 mg), a biphasic microextraction was performed in triplicate following: i) first, 1.5 mL of Methyl-tert butyl ether (MtBE)/MeOH (75:25, v/v), then vortex and sonication for 3 min, ii) second, 1.5 mL of water/MeOH (75:25, v/v), then vortex and sonication for 3 min. The suspension was centrifuged at 14,000 rpm for 3 min at 5°C, then, 500 µL of medium polar extract was taken and dried at 30°C using a vacuum concentrator. The residue was then resuspended using methanol and then stored at -30 °C prior analysis. A quality control (QC) sample was made by pooling aliquots from each extract. Final concentration for untargeted metabolomic analyses was performed at 50-fold dilutions.

On washed samples, an ultra-high-performance liquid chromatography high-resolution mass spectrometry (UHPLC−HRMS) analyses were performed on a Q Exactive™ Plus Hybrid Quadrupole-Orbitrap™ Mass Spectrometer equipped with a Heated Electrospray Ionization (HESI-II) Probe coupled to a UltiMate™ 3000 Rapid Separation LC (RSLC) system (Thermo Fisher Scientific, Villebon-Sur-Yvette, France). Separation was performed on a Luna Omega Polar C18 UHPLC column (150 mm × 2.1 mm i.d., 1.6 μm) with a guard column (Phenomenex, Sartrouville, France). The mobile phase gradient used water containing 0.05 % formic acid (A) and acetonitrile containing 0.05% of formic acid (B). A gradient method at a constant flow rate of 0.3 mL/min was applied under the following conditions: 0 % B for 1 min to 100 % B in 20 min, held at 100 % B over 4 min, then equilibration with 100 % over 4 min. The column oven temperature was set to 40°C, the autosampler temperature was set to 5°C, and the injection volume was fixed to 1 μL. Mass detection was performed in positive ionization (PI) and negative ionization (NI) modes at resolution 30,000 [fullwidth at half-maximum (fwhm) at 400m/z] for MS1 and resolution 17,500 with automatic gain control (AGC) target of 1×10^6^ for full scan MS1 and 1×10^5^ for MS2. Ionization spray voltage was set to 3.5 kV (for PI) and 2.5 kV (for NI), and the capillary temperature was set to 256 °C for both modes. The mass scanning range was *m/z* 100−1500. Each full MS scan was followed by a data-dependent acquisition of MS/MS spectra for the six most intense ions using stepped normalized collision energy of 20, 40, and 60 eV.

#### 2.2.2. Targeted metabolomics

For each grain sample (5 g), extraction was performed in triplicate using 20 mL of ethyl acetate and sonication for 1 h. The suspension was centrifuged at 14,000 rpm for 10 min, and the supernatant dried at 30°C using a vacuum concentrator. The residue was then resuspended using methanol and then stored at -30 °C prior analysis. Final concentration for targeted metabolomic analyses was performed at 12-fold dilutions.

On all set samples in **supp info 1**, a Triple Quad 5500+ LC-MS/MS System – QTRAP Ready (AB Sciex, Villebon sur Yvette, France) equipped with a Turbo Spray source. Separation was performed on an ACQUITY UPLC® BEH C18 column (100 mm × 2.1 mm i.d., 1.7 μm, Waters, MA, USA) equipped with a guard column. The mobile phase gradient used water containing 0.1 % formic acid (A) and acetonitrile containing 0.1% of formic acid (B). A flow rate of 0.3 mL/min was applied following: isocratic conditions, 10% of B and 90% of A for 3 min, then, a gradient of B until 70% in 7 min, and went up to 90% in 1.1 min (curve 6). After 1.9 min, it went back to the initial condition in 0.1 min, and then a re-equilibration for 2.9 min. The column oven temperature was set to 40°C, the autosampler temperature was set to 5°C, and the injection volume was fixed to 5 μL. Mass detection was performed in PI and NI modes at nominal resolution. Ionization spray voltage was set to 4.5 kV (for PI) and 4.5 kV (for NI), and the capillary temperature was set to 500 °C, curtain gas set at 20, ion source gas at 1 at 60 and ion source gas 2 at 70 for both modes. Two main transitions for each standard were performed in PI and NI (details in **supp info 2**). A curve of calibration was carried out for targeted metabolomics from a mix of standards to final concentrations of: 100 ng/mL, 10 ng/mL, 1 ng/mL, 0.1 ng/mL, 0.01 ng/mL, and 0.001 ng/mL in acetonitrile.

### 2.3. Data mining process

#### 2.3.1. Untargeted metabolomics

LC-MS data were processed according to the MSCleanR workflow (Fraisier-Vannier et al., 2020; Vásquez-Ocmín et al., 2021). Briefly, batches in PI and NI were processed separately with MS-DIAL version 4.80 (Tsugawa, Cajka, et al., 2015). MS1 and MS2 tolerances were set to 0.01 and 0.05 Da, respectively, in centroid mode for each dataset. Peaks were aligned on QC reference with an RT tolerance of 0.2 min, a mass tolerance of 0.015 Da, and a minimum peak height detection at 1 × 10^6^ for both ionization modes. MS-DIAL data (both PI and NI) were deconvoluted together with MS-CleanR by selecting all filters with a minimum blank ratio set to 0.8 and a maximum relative standard deviation (RSD) set to 40. The maximum mass difference for feature relationships detection was set to 0.005 Da, and the maximum RT difference was set to 0.025 min. The Pearson correlation links were applied for a correlation ≥ 0.8 and a p-value significance threshold = 0.05. Two peaks were kept in each cluster for further database requests and the kept features were annotated with MS-FINDER version 3.52 (Tsugawa et al., 2016). The MS1 and MS2 tolerances were set to 5 and 15 ppm, respectively. The formula finder was exclusively processed with C, H, O, and N atoms. Two pathways from several sets of data were carried out for the compound’s annotation. For metabolite annotation at level 1: experimental LC-MS/MS data for 500 compounds (including five mycotoxins standards), analyzed on the same instrument and with the same method, were used as references using MS-DIAL based on accurate and exact mass, fragmentation, and retention time. For metabolite annotation level 2: the MS-DIAL, MONA (MassBank of North America), and GNPS (Global Natural Product Social Molecular Networking) mass spectral records were used for spectral match using a dot product score cut-off of 800. The *in silico* match (annotation level 3) were prioritized according to: 1) compounds identified in the literature for both *Chenopodium* and *Amaranthus* genera, and both *Amaranthaceae* and *Chenopodiaceae* botanic family (Dictionary of Natural Products version 28.2, CRC press) mined by MS Finder based on exact mass and in silico fragmentation; 2) generic databases included in MS-FINDER (*i*.*e*., FooDB, PlantCyc, ChEBI, T3DB, Natural Products Atlas, NANPDB, COCONUT, and KNApSAcK); 3)fungal compounds (including mycotoxins) were performed using four “in-house” databases from literature to compile: i) only mycotoxins (n=300) and ii) compounds produced from major mycotoxigenic fungi *Fusarium, Aspergillus*, and *Penicillium*.

Mass spectra similarity networking was carried out from PI mode using MetGem (Olivon et al., 2018) on the final annotated .msp file and metadata files compiled for both batches obtained with MSCleanR. Values used were MS2 *m/z* tolerance = 0.02 Da, minimum matched peaks = 4 and minimal cosine score value = 0.7. Visualization of the molecular network (MN) was performed on Cytoscape version 3.9.1 (Shannon et al., 2003). Original LCMS raw data were uploaded in zenodo (DOI:xxxx)

#### 2.3.2. Targeted metabolomics and mycotoxins quantification

Dataset acquired in format .wiff was converted in .abf format using Analysis Base File Converter (Reifycs, Tokyo, Japan). Analyses were performed with MRMPROBS program version 2.60 (Tsugawa et al., 2013; Tsugawa, Ohta, et al., 2015), using as parameters of peak detection: five scans for minimum peak width; 2,000 as minimum peak height; 0.2 min as retention time tolerance for peak identification; 15% as amplitude tolerance; and 70% as minimum posterior. A list of transitions in .txt format was used as a compound library file. A final list containing the information about the Area above zero for each transition selected in the samples was exported in .txt format. From the final list of information about the mycotoxin standards, a calibration curve was created for each concentration mentioned above. Validation parameters were determined following the International Conference on Harmonization (ICH) Guidelines (ICH, 2005), and the following characteristics were evaluated: 1) linearity, 2) limit of detection (LOD), and 3) limit of quantification (LOQ). Wheat contaminated with enniatins, and beauvericin was used as positive control. Original LC-MS raw data were uploaded in zenodo (DOI:10.5281/zenodo.6700223).

### 2.4. Statistical analyses

The final annotated metabolome dataset generated by the MS-CleanR workflow was uploaded on the MetaboAnalyst 5.0 online platform (Chong et al., 2019). The data were normalized by sum and scaled by Pareto before statistical analysis. First, principal component analysis (PCA) was applied as an exploratory data analysis to provide an overview of LC-MS fingerprints. Then, a hierarchical cluster analysis (HCA) was performed to obtain a dendrogram of varieties using the untargeted metabolome dataset. For each variable the average value across replicates was calculated; then distance matrices were obtained using either 1-r (Pearson correlation coefficient), followed by HCA on each matrix using Ward’s minimum variance criterion using Orange 3.32.0 (Demšar et al., 2013). On the clusters obtained, a partial least squares discriminant analysis (PLS-DA) was conducted using clusters as Y-value, and the top 50 features were plotted on a heatmap using ANOVA. GraphPad Prism 5.0 was used to perform the bar-plot for data quantification.

### 2.5. Fungi isolation and molecular identification

For each sample (**supp info 1**), dried seeds were deposited on Malt Extract Agar (MEA) plates (diameter 15 cm) containing chloramphenicol (100 μg/mL) under the laminar airflow (LAF). Plates were incubated at 27°C daylight. After at least 48h of incubation, growing fungi were replicated on MEA plates without chloramphenicol and incubated at 27°C, with daylight. All the isolated mycelia are stored in UMR 152 laboratory collection (cryopreservation at -80°C in 20% of glycerol).

Total DNA was extracted from mycelium directly picked from the plate, using the Wizard® Genomic DNA Purification kit (Promega, Charbonnières-Les-Bains, France). The final pellet was resuspended in 40 μL of sterile ultra-high quality (UHQ) water and diluted 10 times in UHQ water. The ITS rDNA region was amplified by polymerase chain reaction (PCR) using the primer set ITS5/ITS4 (White et al., 1990). Reactions were carried out in a final volume of 25 μL, containing 2 μL of diluted DNA, 5 µL GoTaq polymerase buffer 5x (Promega), 0.5 μL of each primer, 0.5 µL of dNTP, 0.5 units of GoTaq® Hot Start polymerase (Promega), and 16.5 µL of UHQ sterile water. PCR cycling conditions of the Mastercycler® Nexus Thermal Cycler (Eppendorf AG, Hamburg, Germany) were as follows: initial denaturation at 95°C for 3 min; 35 cycles at 95°C for 45 s, 55°C for 45 s, 72°C for 1 min; and a final extension at 72°C for 10 min. For identification at the species level of the *Aspergillus* and *Penicillium* isolates, beta-tubuline and calmoduline regions were PCR-amplified using the primers Bt2a/Bt2b (Glass & Donaldson, 1995) and CMD5/CMD6 (Hong et al., 2005), in the same conditions as above, except for annealing temperature (58°C instead of 55°C).

PCR amplicons were sequenced by Eurofins Genomics (Ebersberg, Germany) according to a Sanger method using ITS5, Bt2a, and CMD5 respectively. Sequences were submitted to the BLASTn program (http://blast.ncbi.nlm.nih.gov/Blast.cgi) to determine the close matches in the GenBank database. All ITS sequences were aligned with MAFFT v6.814b (Katoh at al 2002) using Geneious®6.1.8. The PhyML method (Guindon & Gascuel, 2003) was used *via* the Geneious® platform to generate a maximum-likelihood phylogenetic tree with the following setting: GTR substitution model, 100 bootstraps, estimated transition/transversion ratio, the estimated proportion of invariable sites, estimated gamma distribution, branch length, and optimized substitution rate. Phylogenetic trees were visualized and edited with MEGAX (Kumar et al., 2018).

## 3. Results and discussion

### 3.1. Untargeted metabolomics for the metabotyping of Andean pseudocereals

Nowadays, chemotype discrimination between plants of the same botanical genus or family represents a challenge in the functional food sector. Chemotypes are characterized by the presence of certain metabolites, as major or minor compounds, in a plant. Despite their potential to enhance the food basket, the chemodiversity of pseudocereals has received little attention so far. To explore this chemodiversity, we used an untargeted metabolomics approach referred to as “metabotyping” as we did in a previous work investigating Cannabis cultivars (Vásquez-Ocmín et al., 2021). We thus analyzed the whole metabolomes of several pseudocereal grains collected in the Andes of Peru, as well as one quinoa sample from France (**Fig. 1-A**). To ensure a comprehensive analysis of medium polar extract, positive ionization mode was investigated using the MS-CleanR workflow, which resulted in a list of 219 features (*m/z* x RT). Depending on the level of annotation mentioned above, 19 compounds were yielded for the genus (8.7%), 43 compounds for the family (19.6%), 128 compounds for the generic level (58.4 %), and 29 unknowns (13.2 %). The fitness of compound annotation generally depends on the databases (DB) used. Despite the few studies on pseudocereals, we confirmed an intern DB from family and genera gathered from the literature, responsible for 28.3 % of annotations (annotation level 3-1). This was complemented with our experimental DB (annotation level 1) and the spectral match with other DB in line (annotation level 2). Using NPClassifire and ClassyFire (Djoumbou Feunang et al., 2016; Kim et al., 2021) the major classes and subclasses of compounds following their biosynthetic pathway were identified (**supp info 3**).

**Figure 1.**
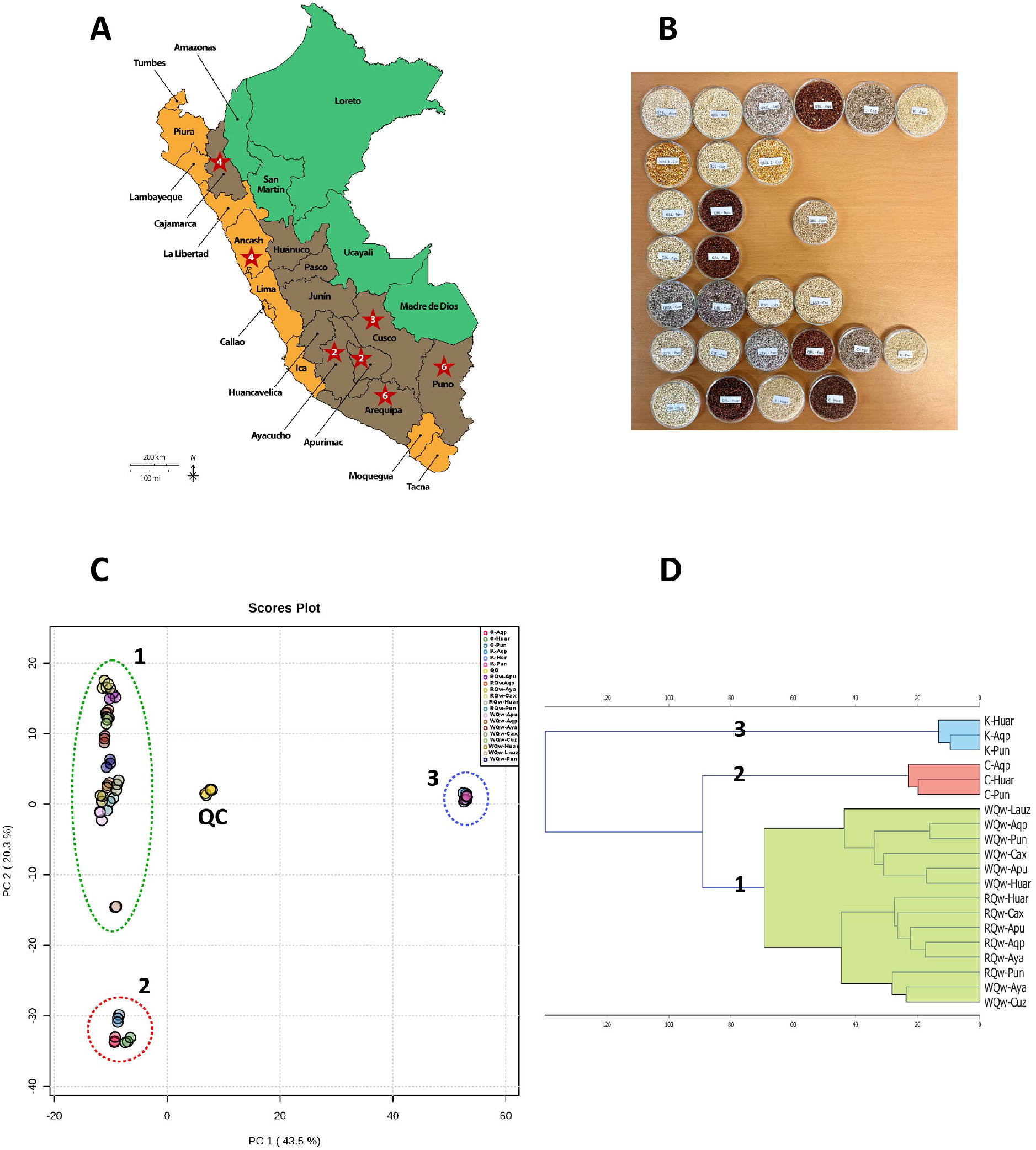
(**A**) Geographic origins of the grain samples. The names of the 24 administrative departments of Peru are indicated in black. The departments of collection are stamped with a red star marked with the number of samples collected therein. Green = Peruvian Amazon (including highland and lowland jungles); brown = Andean highlands; orange = arid and semi-arid Pacific coastal lands. (**B**) Set of samples used for the metabolomics analyses. (**C**) Global PCA of 20 pseudocereal grains from Peru (except for WQ-Lauz, France). (**D**) Hierarchical clustering analysis (HCA) following the whole metabolome to group all grain samples. Pseudocereal varieties: WQw-Cuz = white quinoa washed from Cuzco, WQw-Aya = white quinoa washed from Ayacucho, RQw-Aya = red quinoa washed from Ayacucho, WQw-Apu = white quinoa washed from Apurimac, RQw-Apu = red quinoa washed from Apurimac, WQw-Cax = white quinoa washed from Cajamarca, RQw-Cax = red quinoa washed from Cajamarca, WQw-Pun = white quinoa washed from Puno, RQw-Pun = red quinoa washed from Puno, K-Pun = kiwicha from Puno, C-Pun = canihua from Puno, WQw-Aqp = white quinoa washed from Arequipa, RQw-Aqp = red quinoa washed from Arequipa, K-Aqp = kiwicha from Arequipa, C-Aqp = canihua from Arequipa, WQw-Huar = white quinoa washed from Huaraz, RQw-Huar = red quinoa washed from Huaraz, K-Huar = kiwicha from Huaraz, C-Huar = canihua from Huaraz, WQw-Lauz = white quinoa washed from Lauzerte, France.

Phylogenetic taxonomy has grouped the previously separated subfamily *Chenopodiaceae* within the family *Amaranthaceae* s.l., which include now the five phylotranscriptomic clades *Amaranthaceae* s.s., *Betoideae*, “Chenopods I”, “Chenopods II”, and *Polycnemoideae* (Morales-Briones et al., 2020). In our study, we have profiled by LC-MS 20 pseudocereal grain samples that belonged to: Chenopods I [*Chenopodium quinoa* (14 samples) and *C. pallidicaule* (3 samples)] and *Amaranthaceae* s.s. [*Amaranthus caudatus* (*syn*: *A. cruentus*) (3 samples)]. To obtain an unsupervised overview of chemical fingerprint distribution, we firstly performed a principal component analysis (PCA). The PCA score plot from the two first principal components accounted for 63.8% of the total variance (**Fig. 1- B**). As expected, repeated injections of QC were grouped near the center of the PCA score plot demonstrating a good reproducibility and stability of the analytical workflow. Then, a hierarchical clustering analysis (HCA) was performed with 20 samples (three analytical repeats for each sample), obtaining three very well differenced clusters (**Fig. 1-C**). So, our metabolomic signature showed that *Chenopodium* species are grouped in two distinct clusters: **cluster 1** (14 samples of **quinoa**) and **cluster 2** (3 samples of **canihua**). **Cluster 3** brought together the three *Amaranthus* grains (**kiwicha**). On theother hand, the provenance (in all clusters) and color of grains (in the **cluster quinoa**) did not play a significant role in the metabolomic classification intraspecies. However, inside the **cluster quinoa**, two other subgroups for red and white quinoa can be observed. An exception is white quinoa samples from Ayacucho and Cuzco that were compiled with red quinoa samples, which could infer a putative hybridization intraspecies. Additionally, to validate the HCA model and highlight characteristic biomarkers of each cluster, we performed a supervised Partial Least Squares-Discriminant Analysis (PLS-DA) based on the variable importance in projection (VIP) and a heat map was generated from the top 50 metabolites selected by Anova (**supp info 4, 6-8**).

After our workflow analysis, unanticipated metabolites produced by filamentous fungi were identified in the final metadata. These metabolites were placed in a molecular network (**Fig. 2-A**). The molecular network of the whole dataset analyzed in PI mode was constructed to identify possible similarities between molecules based on spectral mass similarity (MS-MS fragmentation patterns) (**Fig. 2**). By grouping the major classes of compounds in clusters according to their biosynthetic pathway, there were a number of characteristic groups, such as amino acids and peptides, carbohydrates, alkaloids, fatty acids, polyketides, shikimates and phenylpropanoids, terpenoids, and others. Here, we can also observe the macrolide derivatives pandangolide 1a isolated from *Cladosporium sp*. (Gesner et al., 2005), and chaxlactin A from *Streptomyces* strain (Rateb et al., 2011), the indole diketopiperazine alkaloid derivative asperfumifatin previously isolated from the endophytic *Aspergillus fumigatus* (Xie et al., 2015), and the polyketide derivative altersolanol O from *Alternaria sp*. (Chen et al., 2014) (**Fig. 2-A**). Another compound identified was a zearalenone derivative (3-(14,16-dihydroxy-3-methyl-1,7- dioxo-3,4,5,6,7,8,9,10-octahydro-1H-2-benzoxacyclotetradecin-15-yl)-N-[2-(1H-imidazol-1-yl)ethyl]-3- (3,4,5-trimethoxyphenyl) propenamide). This compound was also identified as a marker for the **cluster kiwicha** (**supp info 4-C**). Zearalenone (ZEA) type of compounds are produced by several mycotoxigenic *Fusarium* (*F. graminearum, F. cerealis, F. culmorum, F. equiseti*, etc.), and are considered as mycotoxins. ZEA is distributed worldwide and naturally occurs in agricultural crops, particularly in maize. This mycotoxin could contaminate products made of barley, wheat, oats, rice, sorghum, and others (G.-L. Zhang et al., 2018). The structure is a macrolide type, and the name “zearalenone” is derived from the combination of the terms maize (Zea mays)—“zea”, resorcylic acid lactone—“ral”, — “en” for the presence of a double-bond, and “one” for the ketone group (Ropejko & Twarużek, 2021). The flexibility of the molecule is enough to adopt a conformation able to bind to the mammalian estrogen receptor, although with an affinity lower than the natural estrogen 17-β-estradiol resulting in severe effects on the reproductive system in several animal species, particularly pigs (Galaverna & Dall’Asta, 2012; G.-L. Zhang et al., 2018). Even if the molecule identified has the same macrolide structure of ZEA, this latter also presents a tyrosine alkaloid moiety. So far, to our knowledge, no bibliography about their toxicity was found. In this sense, close attention was taken to the compounds having structural similarities. In addition, another macrolide derivative (retrorsine) previously reported from *Senecio gilliesiano* or *S. gilliesianus* (Asteraceae) (Guidugli et al., 1986) was annotated. All these metabolites described above are placed as self-loop in the molecular network. This suggests that there are no other molecule derivatives in the whole metabolome of the grains.

**Figure 2.**
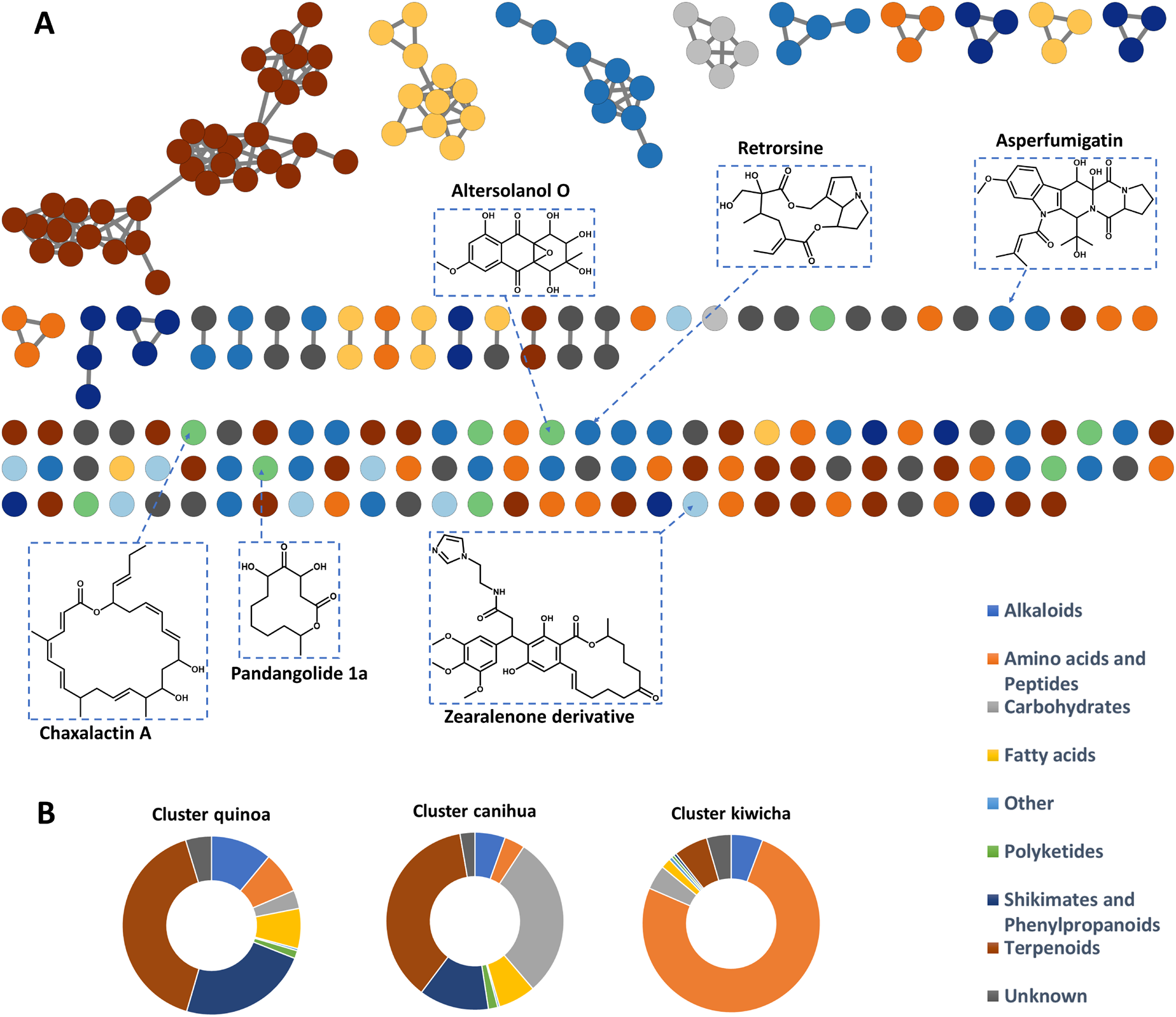
Molecular network (PI mode) from compounds identified in pseudocereals, based on MetGem and visualized by Cytoscape. Colors in **A** and **B** are correlated with the major classes of compounds annotated. Structures and names of some metabolites annotated from fungal biosources are represented in **A**. In **B**, the clusters are correlated with **Fig 1-C**. In these clusters, we represented the percentage of the major classes of compounds obtained by cluster.

Moreover, we displayed the information correlated between the major class of compounds and the cluster producing (**Fig. 2-B**). As described in the literature, here we can observe that terpenoids (41%) and molecules belonging at shikimates and phenylpropanoids (24%) (*e*.*g*. flavonols, anthocyanidins, cinnamic acid derivatives, etc) are the most representative for the **cluster quinoa** (Lin et al., 2019; Tabatabaei et al., 2021). Among the terpenoids, saponins were not identified, it is due to the postharvest process, where the grains are washed on various days with plenty of water to remove saponins localized in the pericarp (Jarvis et al., 2017). Triterpene saponins are known for their ecological defense role in the plant but are also known by the local people for providing a bitterness “*amargor*” or bitter-tasting “*sabor amargo*” to the unwashed grains (McCartney et al., 2019). For the **cluster canihua**, terpenoids (37%) and carbohydrates (29%) are the most representative class of molecules. At the opposite of the *Chenopodium* grains of the **clusters quinoa** and **canihua**, the majoritarian compounds identified for the **cluster kiwicha** were the amino acids (76%), mainly small peptides (**Fig. 2-B**). Due to the nutritional richness, in the last years the pseudocereals studied here are appreciated worldwide (Fletcher, 2016; Perez-Rea & Antezana-Gomez, 2018).

### 3.2. Targeted metabolomics revels the presence of an emerging mycotoxin in Andean pseudocereals

We complemented our approach with a targeted metabolomic analysis using LC-MS Qtrap Multiple Reaction Monitoring (Qtrap-MRM). Specifically, we focused on Aflatoxin B1 (AflaB1), Beauvericin (BEA), Fumonisin B1 (FumB1), Ochratoxin (OTA), and Patulin (Pat), which are mycotoxins produced by several species of *Fusarium, Aspergillus*, and *Penicillium*. Before implementing QqQ- MRM, we designed the main transitions (fragments) in both PI and NI modes of mycotoxins standards, as described previously (**sup. info 2**) (Stead et al., 2014). We then assessed whether post-harvest washing could eliminate mycotoxins by comparing targeted metabolomes of washed and unwashed quinoa grains. On the five mycotoxins targeted, only FumB_1_ and BEA were detected in PI mode (**supp info 9**). Among all samples, only seven of the quinoa samples trace amounts of FumB_1_ (not quantifiable, <LOQ) (**supp info 10**). This result supports the interest in the consumption of quinoa as safe food, taking into account that our team has previously reported that other Andean grains may be contaminated with high levels of FumB_1_ (notably maizes), exceeding the maximum food safety level of 1,000 ng/g (Ducos et al., 2021). Fumonisins, which have a long carbon chain aminoalcohol structure as their basic skeleton, are structurally similar to sphingosines (sphingoids). These mycotoxins are primarily produced by *F. verticillioides* and *F. proliferatum* (Smith, 2018). It has been suggested that FumB_1_ may increase the risk of cancer, notably in kidney and liver. Thus, FumB_1_ was classified by the International Agency for Research on Cancer (IARC) as a potential human carcinogen (Group 2B) (Claeys et al., 2020; Ostry et al., 2017). Concerning BEA, it was detected in 17 pseudocereal grain samples from Peru and in the only sample from France, with levels equal to or greater than the limit reported by the European Food Safety Authority (EFSA) for dietary acute exposure (*i*.*e*., 0.05 ng/mL or 0.14 ng/g of dried seeds) (**Fig. 3-A**) (EFSA, 2014). BEA and enniatins are emerging mycotoxins since they are not routinely determined nor regulated like other *Fusarium* mycotoxins, but evidence of their occurrence is rapidly increasing as an important natural contaminant in cereals and cereal-derived foods (Malachova et al., 2011; Vaclavikova et al., 2013). BEA is a cyclic hexadepsipeptide synthesized by mycotoxigenic fungi, such as *Beauveria bassiana* and *Fusarium* spp. (Wu et al., 2018). This mycotoxin is harmful to human cells, displaying toxicity at lower concentrations than AflaB_1_ (Svingen et al., 2017). Disturbingly, our findings indicate very elevated levels of BEA (7.9 ± 0.85 ng/g) in washed quinoa samples from Cajabamba (region of Cajamarca), 150-fold higher than the limit set by EFSA. Cajabamba is located on the eastern slopes of the Andes (2,648 meters height above mean sea level, toward the highland jungle); hence, its ecoregion might have favorable conditions for the formation of mycotoxins by fungi (temperature, pH, moisture, etc.). The contamination of quinoa grains with BEA originating from Cajamarca represents a point of concern regarding food safety, as local authorities have recently promoted the cultivation of quinoa in this region (Gobierno Regional de Cajamarca, 2020). Significant amounts of BEA were also detected in canihua samples from Puno (C-Pun), with quinoa samples from France (WQw-Lauz) and red quinoa samples from Arequipa (RQw-Aqp). We also compared the relative mean peak area of detected mycotoxin in the untargeted metabolomics approach. In this sense, asperfumigatin, and altersolanol O are more present in samples for the quinoa and zearalenone derivative for all kiwicha samples (**Fig 3-B**). In **cluster quinoa**, the samples with more presence of asperfumigatin were red quinoa from Huaraz, Arequipa and Cajamarca, and white quinoa from Cajamarca (RQw-Huar, RQw-Aqp, RQw-Cax, WQw-Cax); and for altersolanol O, mainly white quinoa from Arequipa and Ayacucho, and red quinoa from Puno (WQw-Aqp, and WQw-Aya, RQw-Pun) (data not showed).

**Figure 3.**
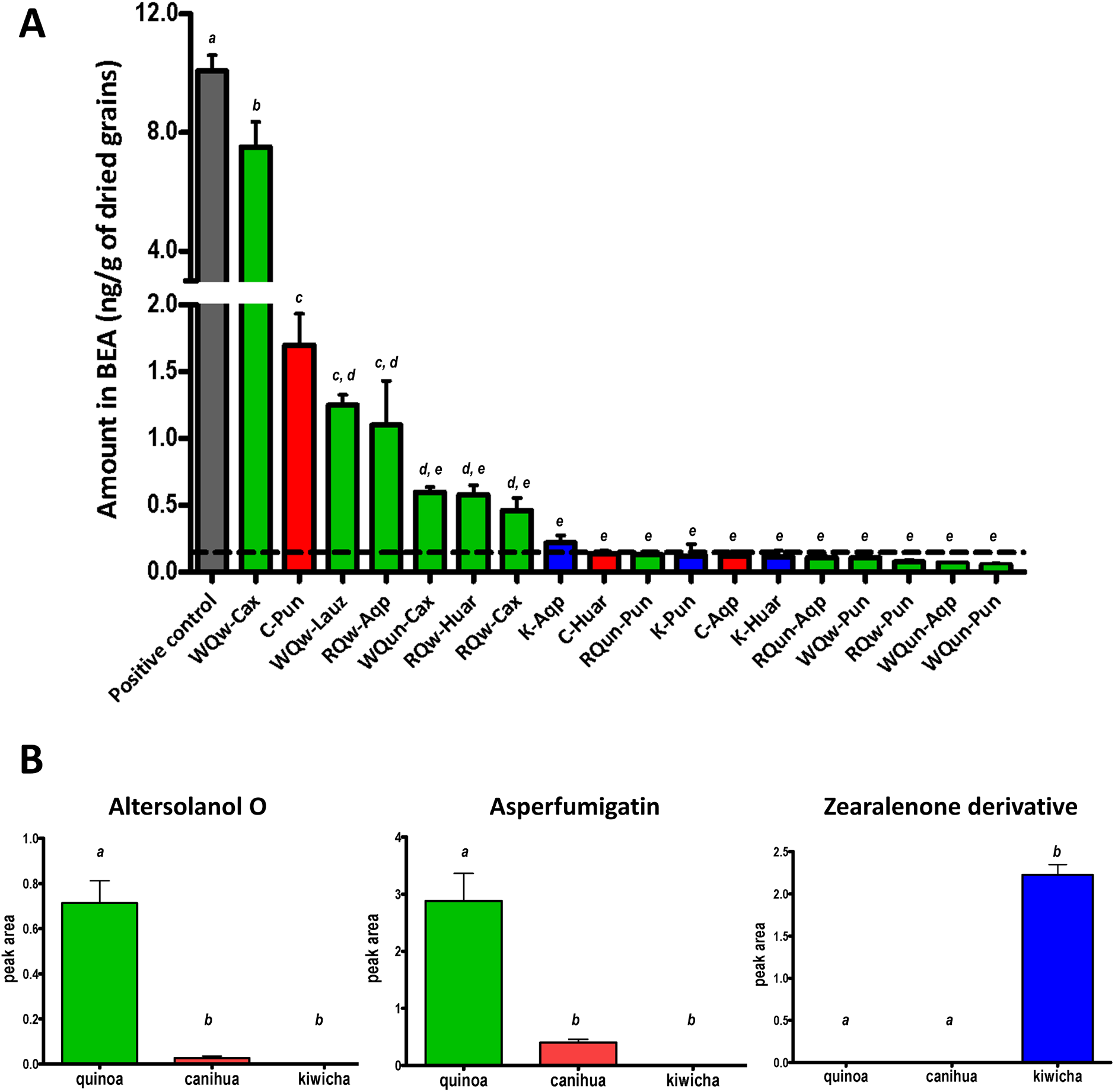
Quantification of BEA mycotoxin and others metabolites by LCMS. (**A**) Bar-plot of BEA amount (mean + sd, n=3) in Andean pseudocereal samples expressed in ng/g of dried grains. The dotted line represents the theoretical level of beauvericin among the legislation (0.14 ng/g in dried seeds, EFSA). (**B**) Bar-plot displaying the relative quantification by the cluster of three molecules produced by fungus microorganisms in AP. The lowercase letters indicate significant differences between pseudocereal samples (one-way ANOVA).

### 3.3. Fungal inventory infesting the Andean pseudocereals

The fungal inventory was realized with the aims to better know the filamentous fungi presents in the Andean pseudocereal samples and try to correlate this information with the presence of molecules produced themselves, especially the mycotoxins. Infestation by at least one fungal endobiont was spotted in 22 up 28 grain samples (78%). Sequence analysis was able to identify 74 fungal isolates (out of 85 total) at the genus level. Overall, we pinpointed 15 fungal families, composed by 20 genera and 14 species (**supp info 11**). Ascomycetes were the most common phylum of fungi (84%) (**Fig. 4 A-B**), represented by five orders: i) Eurotioales (*Aspergillus* sp., *Penicillium* sp., and *Talaromycetes* sp.), represents nearly two-third of the strains isolated (58%); ii) Pleosporales (19%), comprising: *Alternaria* sp., *Bipolaris* sp., *Didymella* sp., and *Epicoccum* sp.; iii) Cladosporales (11%) (only *Cladosporium* sp.); iv) Hypocreales (6%), and v) Xylariales (6%). In the same way, five Basidiomycetes and two Mucoromycetes species were spotted. Based on the ITS phylogeny and beta-tubulin and calmoduline genes analysis, a large diversity of *Aspergillus* sp. and *Penicillium* species could be observed. However, the precise identification of some others species was not achieved using ITS5 primers and the utilization of genus specifics primers (*i*.*e*. gpd 1 for *Alternaria*) would allow to go further in identifications (Berbee et al., 1999).

**Figure 4.**
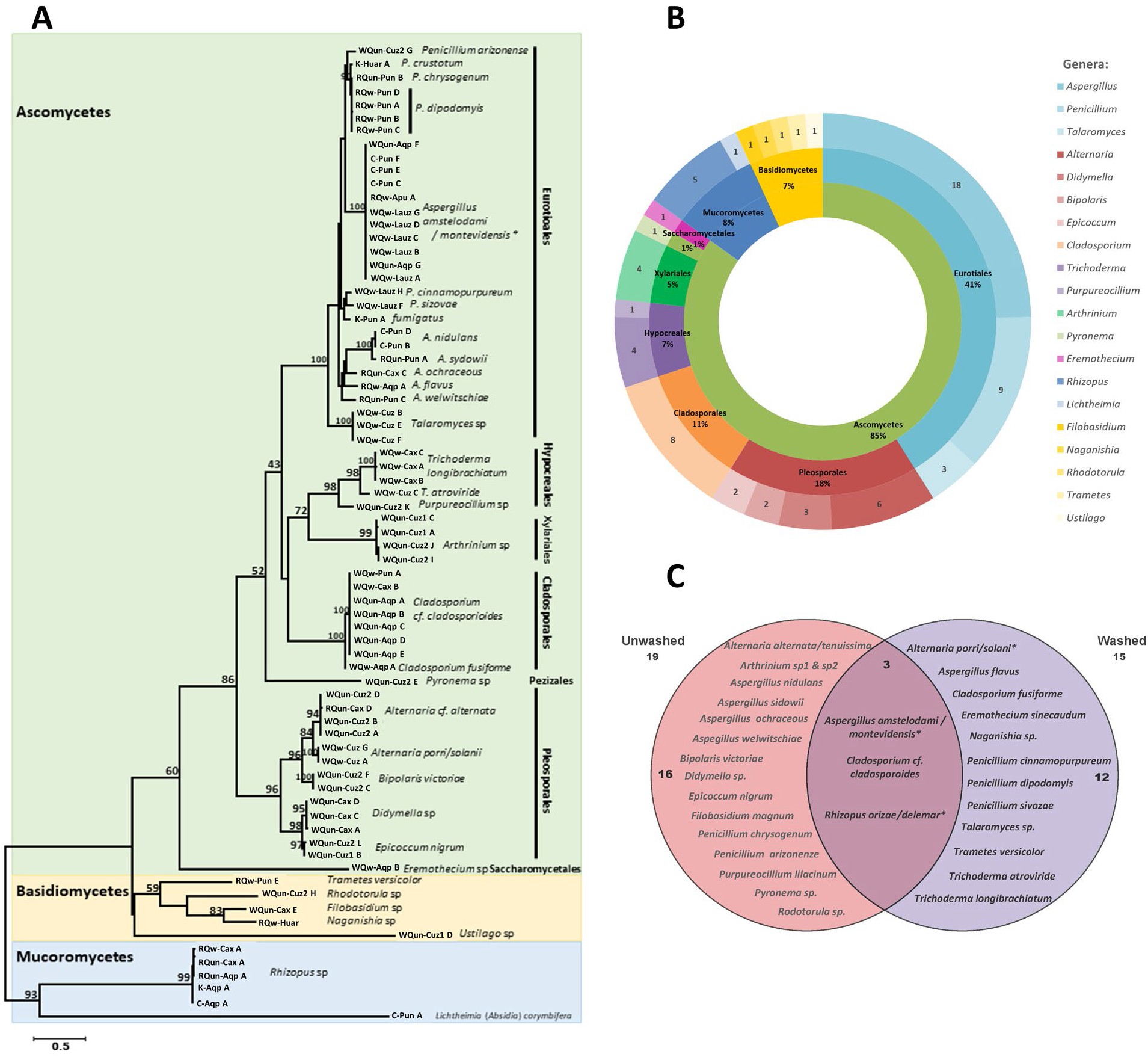
Maximum-likelihood phylogenetic tree based on samples ITS sequences and closest ITS reference sequences from GenBank. The tree was obtained by applying the PhyML method in the Geneious® plateform. Bootstrap values > 50% are indicated above branches. (**A**) Codes of samples are described in **supp info 1** and letters were assigned for each strain isolated. (**B**) Main genus of fungal community identified from Andean pseudocereals, genera are expressed in number of strains and percentage. (**C**) Diagram of Venn for washed and unwashed conditions from quinoa relied to their fungal community. * : not possible to choose between the 2 species

Mycotoxigenic fungi have been traditionally classified into two main groups: i) field fungi that contaminate crops before and during harvest (*e*.*g*., *Fusarium, Alternaria*, and *Cladosporium* spp.) and, ii) storage fungi (e.g., *Penicillium* and *Aspergillus* spp.) that tend to contaminate crops in a post-harvest manner. Overall, the mycotoxigenic fungi we found attached to Andean pseudocereals belonged to several genera, including *Aspergillus, Penicillium*, and *Alternaria*. For instance, *Alternaria* sp. is an ubiquitous plant pathogen that can infest a broad spectrum of plants, causing damage to crops and representing a hazard to human health (Troncoso-Rojas & Tiznado-Hernández, 2014). Moreover, some genera have also the ability to protect their host against phytopathogens, such as the heterogeneous genus *Cladosporium* able to synthesize metabolites such as cladosporine with antifungal activity on the spore germination of *Penicillium* and *Aspergillus* species (Bensch et al., 2015; Scott et al., 1971).

In our study even if BEA was detected in several samples, our fungal inventory about pseudocereals did not show the presence of *Fusaria* or *Beauveria*. It could be explained by fact that mycotoxin may persist while the fungus has disappeared. In addition, the natural production of these fungal groups is very sensitive to many environmental factors, *i*.*e*., altitude, drought, etc. Experimental limits, mainly the use of a limited number of seeds for each sample leading to non-exhaustive fungal isolation could also explain the absence of a *Fusarium* community in our inventory. Indeed, in our experimental condition, others species, better adapted to our experimental design might inhibit the growth of other strains. Likewise, the presence of the fungus does not prove the presence of mycotoxins. Some species are kwon to inhibit the mycotoxins production of other species such as the post-harvest pathogen *Rhizopus oryzae*, able to degrade mycotoxins like aflatoxins, OTA, or patulin, by the presence of a specific enzyme (Varga et al., 2005). The strain *Rhizopus sp*. was present concomitantly in our samples (RQw-Aqp and RQun-Cax) infested by *A. flavus* and *A. cf. ochraceus*, producers of aflatoxins like AflaB_1_ or OTA. Our chemical analysis did not detect AflaB_1_ and OTA in these samples, so we hypothesized that *Rhizopus sp*. could have detoxified the sample of the mycotoxins. On the other hand, our inventory highlighted the presence of *Alternaria, Aspergillus*, and *Cladosporium* which could validate the metabolites production of alternolanol O (*Alternaria sp*.), asperfumigatin (*Aspergillus fumigatus*), and Pandangolide 1a (*Cladosporium sp*.), respectively.

A comparison between fungal species encountered on washed and unwashed quinoa was also performed. Nineteen species were isolated on unwashed seeds and 11 on washed samples (**Fig 4-C**). Among them, only three species grew in both conditions: *Aspergillus amstelodami/montevidensis, Cladosporium cf. cladosporoides*, and *Rhizopus sp*. These washing steps have double consequences: on the one hand, it can cause the elimination of the fungal strains present on seeds at the time of the harvest, and, on the other hand, this phenomenon involving water can allow the emergence of other strains being able to develop under favorable conditions.

## 4. Conclusion

Global interest in pseudocereals as “superfood” has increased over the last decade. However, their chemodiversity has not been well studied to date. In the present work, metabotyping was able to add new insights into the chemotaxonomy of pseudocereals, confirming the previously established phylotranscriptomic clades. Metabolomics identified three clusters among *Chenopodium* (**clusters 1** and **2**) and *Amaranthus* grain samples (**cluster 3**). In addition, we highlighted their main markers and major class of compounds by cluster. Moreover, we report for the first time the presence of mycotoxins in the pseudocereals. Sixteen samples of Peru (out of 27) and one sample from France (out of one) were contaminated with Beauvericin, an emerging mycotoxin. One sample of white quinoa from Cajabamba (Cajamarca) showed an alarming high level of Beauvericin. To link the occurrence of mycotoxins with the presence of toxin-producing fungi in pseudocereals, an inventory of filamentous fungi was taken. Several mycotoxigenic fungi were detected, including *Aspergillus sp*., *Penicillium sp*., and *Alternaria sp*., but not *Fusaria*. The present study underscores the need to better understand the chemistry of pseudocereals, as well as further research into the risk associated with emerging mycotoxins.

## Supporting information

Supporting informations

## Authors contribution

Conceptualization: SB, PGV-O, and GM; methodology: PGV-O, GM, and SB; chemical data acquisition/curation: PGVO and GM; biological data curation: AG, PJ, and SB; investigation: PGV-O, GM, AG, SB, GC, JAV-B, SC-Z, PJ, NP, and MH. PGV-O wrote the original draft and all the authors contributed and validated the final manuscript.

## Acknowledges

The authors wish to acknowledge a postdoctoral fellowship of PGV-O from the French National Research Institute for Sustainable Development (IRD) under contract number 04077858. The authors are grateful to Pascal Pineau (Institut Pasteur de Paris), Olivier Puel, Sophie Lorber-Pascal (Toxalim, INRAE) for the provision of mycotoxin standards. The authors are also grateful with Lina Abdelwahed (Université Paul Sabatier – Toulouse 3), Sylvie Fournier, Amélie Pérez, Justine Chervin, Virginie Puech-Pagès (MetaToul-Agromix), Juan Pablo Cerapio Arroyo (Centre de Recherches en Cancérologie de Toulouse), David Condori Cruz (Universidad Peruana Cayetano Heredia) for their technical supports; and Alexandre Maciuk (Université Paris-Saclay) for his critical discussion.

