## Supporting informations for "Metabotyping of Andean pseudocereals and characterization of emerging mycotoxins"

**Table of Content**


1. Set samples and description of pseudocereal grains **Supp info 1**
2. MRM transitions in positive and negative ionizations mode for five mycotoxins studied in this work **Supp info 2**
3. Metadata **Supp info 3**
4. Additional statistical analysis **Supp info 4**
5. Markers by cluster **Supp info 5**
6. Heat map to cluster quinoa (cluster 1) **Supp info 6**
7. Heat map to cluster canihua (cluster 2) **Supp info 7**
8. Heat map to cluster canihua (cluster 3) **Supp info 8**
9. MS-MS pathway fragmentations for BEA and FumB1 by MRM **Supp info 9**
10. Mycotoxins quantification from pseudocereal grains by MRM **Supp info 10**
11. Fungal strains isolated from Andean pseudocereal seeds **Supp info 11**
12. References

**1.** Set samples and description of pseudocereal grains

| Codes/ID | Name | Drying | Cultivation type | Locally, Region, country | Elevation (m) | Comments |
|---|---|---|---|---|---|---|
| **WQun-Cuz1** | White quinoa-unwashed | Open field | Open field | Chumbivilcas, Cuzco, Peru | 3642 | Obtained from a farmer's fair |
| **WQun-Cuz2** | White quinoa-unwashed | Open field | Open field | Quillabamba, Cuzco, Peru | 1 050 | Obtained from a farmer's fair |
| **WQw -Cuz** | White quinoa-washed | Open field | Open field | Quillabamba, Cuzco, Peru | 1 050 | Obtained from a farmer's fair |
| **WQw-Aya** | White quinoa-washed | Open field | Open field | Vilcashuaman, Ayacucho, Peru | 3490 | Obtained from a farmer's fair |
| **RQw-Aya** | Red quinoa-washed | Open field | Open field | Vilcashuaman, Ayacucho, Peru | 3490 | Obtained from a farmer's fair |
| **WQw -Apu** | White quinoa-washed | Open field | Open field | San Jeronimo, Apurimac, Peru | 2954 | Obtained directly from a farm |
| **RQw-Apu** | Red quinoa-washed | Open field | Open field | San Jeronimo, Apurimac, Peru | 2954 | Obtained directly from a farm |
| **WQun-Cax** | White quinoa-unwashed | Open field | Open field | Cajabamba, Cajamarca, Peru | 2648 | Obtained directly from a farm |
| **WQw -Cax** | White quinoa-washed | Open field | Open field | Cajabamba, Cajamarca, Peru | 2648 | Obtained directly from a farm |
| **RQun-Cax** | Red quinoa-unwashed | Open field | Open field | Cajabamba, Cajamarca, Peru | 2648 | Obtained directly from a farm |
| **RQw-Cax** | Red quinoa-washed | Open field | Open field | Cajabamba, Cajamarca, Peru | 2648 | Obtained directly from a farm |
| **WQun-Pun** | White quinoa-unwashed | Open field | Open field | San Roman, Puno, Peru | 3855 | Obtained from a farmer's Coop |
| **WQw-Pun** | White quinoa-washed | Open field | Open field | San Roman, Puno, Peru | 3855 | Obtained from a farmer's Coop |
| **RQun Pun** | Red quinoa-unwashed | Open field | Open field | San Roman, Puno, Peru | 3855 | Obtained from a farmer's Coop |

| | | | | | | |
|---|---|---|---|---|---|---|
| **RQw-Pun** | Red quinoa-washed | Open field | Open field | San Roman, Puno, Peru | 3855 | Obtained from a farmer’s Coop |
| **K-Pun** | Kiwicha | Open field | Open field | San Roman, Puno, Peru | 3855 | Obtained from a farmer’s Coop |
| **C-Pun** | Canihua | Open field | Open field | San Roman, Puno, Peru | 3855 | Obtained from a farmer’s Coop |
| **WQun-Aqp** | White quinoa-unwashed | Mechanical dryer | Open field | Majes, Arequipa, Peru | 1410 | Obtained directly from a farm |
| **WQw-Aqp** | White quinoa-washed | Mechanical dryer | Open field | Majes, Arequipa, Peru | 1410 | Obtained directly from a farm |
| **RQun-Aqp** | Red quinoa-unwashed | Mechanical dryer | Open field | Majes, Arequipa, Peru | 1410 | Obtained directly from a farm |
| **RQw-Aqp** | Red quinoa-washed | Mechanical dryer | Open field | Majes, Arequipa, Peru | 1410 | Obtained directly from a farm |
| **K-Aqp** | Kiwicha | Open field | Open field | Cotahuasi, Arequipa, Peru | 2675 | Obtained from a farmer’s fair |
| **C-Aqp** | Canihua | Open field | Open field | Cotahuasi, Arequipa, Peru | 2675 | Obtained from a farmer’s fair |
| **WQw-Huar** | White quinoa-washed | | | Yungay, Huaraz, Peru | 3038 | Obtained directly from a farm |
| **RQw-Huar** | Red quinoa-washed | | | Yungay, Huaraz, Peru | 3038 | Obtained directly from a farm |
| **K-Huar** | Kiwicha | | | Yungay, Huaraz, Peru | 3038 | Obtained directly from a farm |
| **C-Huar** | Canihua | | | Yungay, Huaraz, Peru | 3038 | Obtained directly from a farm |
| **WQw-Lauz** | White quinoa-washed | | | Lauzerte (82), Occitanie, France | 98 | Obtained in the city market of Toulouse, France |

2. MRM transitions in positive and negative ionizations mode for five mycotoxins studied in this work

| Mycotoxin | MRM transition | RT | Cone voltage (V) | Collision energy (eV) | Polarity |
|---|---|---|---|---|---|
| **Aflatoxin $B_1$** | Q 313.014 > 285.092 | 9.33 | 111 | 31 | Pos |
| | q 313.014 > 240.942 | 9.33 | 111 | 55 | Pos |
| **Beauvericin** | Q 784.374 > 134.111 | 13.48 | 101 | 39 | Pos |
| | q 784.374 > 244.156 | 13.48 | 101 | 77 | Pos |
| **Fumonisin $B_1$** | Q 722.296 > 334.307 | 9.27 | 116 | 55 | Pos |
| | q 722.296 > 352.329 | 9.27 | 116 | 53 | Pos |
| **Ochratoxin A** | Q 403.993 > 239.051 | 11.34 | 26 | 31 | Pos |
| | q 403.993 > 213.100 | 11.34 | 26 | 23 | Pos |
| **Patulin** | Q 154.676 > 72.930 | 1.53 | 86 | 25 | Pos |
| | q 154.676 > 52.984 | 1.53 | 86 | 29 | Pos |
| **Beauvericin** | Q 782.319 > 260.137 | 13.48 | 165 | 30 | Neg |
| | q 782.319 > 168.017 | 13.48 | 165 | 62 | Neg |
| **Fumonisin $B_1$** | Q 720.266 > 156.916 | 9.27 | 30 | 48 | Neg |
| | q 720.266 > 562.407 | 9.27 | 30 | 40 | Neg |
| **Ochratoxin A** | Q 402.156 > 61.997 | 11.34 | 10 | 50 | Neg |
| | q 402.156 > 339.214 | 11.34 | 10 | 12 | Neg |
| **Patulin** | Q 152.942 > 108.997 | 1.53 | 15 | 14 | Neg |
| | q 152.942 > 81.033 | 1.53 | 15 | 14 | Neg |

* Aflatoxin B1 was not detected in ESI(-).

### 3. Metadata

Metadata is available online in zenodo (DOI: 10.5281/zenodo.6700223).

### 4. Additionally statistical analysis

To validate the HCA model and highlight characteristic biomarkers of each cluster, we performed a supervised Partial Least Squares Discriminant Analysis (PLSDA) based on the variable importance in projection (VIP) values (**A-B**). As a result, 63.6% of the total variance was displayed on the first two principal component axes of the PLSDA score plot. The PLSDA confirmed the three clusters previously identified. Among all grain samples, 28 compounds were identified as discriminating metabolites (VIP > 1). For the **clusters quinoa** and **canihua**, as expected main markers are mainly terpenoids and flavonoids regularly presents in the *Chenopodium* genus (Pereira et al., 2019; Rastrelli et al., 1996). From **quinoa**, the triterpenoid garsubellin B isolated from *Garcinia subelliptica* (Fukuyama et al., 1998) and a hypothesis of structure colossolactone III derivative, a triterpenoid of withanolides type, were annotated (Dine et al., 2008). Withanolides are the structures previously isolated from the Solanaceae family (i.e. *Physalis longifolia*, *Vassobia breviflora*, and *Withania somnifera*), with anti-proliferative activity against an array of cell lines (Sahai, 1985; Zhang et al., 2012). Another triterpenoid annotated was the cycloartane tubiferolide methyl ester, previously reported from *Gardenia sootepensis* (Rubiaceae) (Nuanyai et al., 2009). Among the flavonoids, molecules reported from the *Chenopodium* genus were annotated, like kaempferol-3-O-galactoside derivative (*C. murale*), and isorhamnetin 3-(2G-apiosylrutinoside) (*C. pallidicaule*) (Harborne & Baxter, 1999). The only marker identified for **canihua** was the ecdysteroids derivative hexahydroxycholest-7-en-6-one (Inokosterone), previously isolated from the family Amaranthaceae (Harborne & Baxter, 1999). For **kiwicha**, main markers identified were the small amino acid derivative valine betaine, and nucleotide bases like adenosine isomers previously isolated from *Beta vulgaris* (Amaranthaceae) (Poulton & Butt, 1976).

Then, a heat map was generated from the top 50 metabolites (**C**) selected by Anova to find other characteristic markers differentiating these clusters (all the markers by cluster are described in **supp. info 5**). In **quinoa**, only four molecules were discriminant: an isomer of colossolactone III derivative (also identified in the PLSDA-VIP), an unknown compound, an alkaloid derivative (methyl 5-hydroxyoxindole-3-acetate), and a hypothesis of structure identified as an indole alkaloid derivative (leptoclinidamine B) previously isolated from a Australian ascidian Leptoclinides (Carroll & Avery, 2009). In **canihua**, 13 markers were identified, among which the flavonoids previously isolated from the family Chenopodiaceae and Amaranthaceae families: isorhamnetin-3-O-rutinoside from *Salsola kali*, quercetin 3-O-rutinoside previously from *Amaranthus spinosus*, and isorhamnetin 3-O-robinobioside from *Aerva javanica* (Harborne & Baxter, 1999; Hegnauer, 1989). Other compounds identified were the terpenoids pubesenolide previously isolated from *Physalis peruviana* and *P.*

*pubescens* (Solanaceae) (Sahai, 1985) and 2,3,14,20,22,26-Hexahydroxycholest-7-en-6-one previously isolated from *Achyranthes bidentata* (Amaranthaceae). In **kiwicha**, 33 molecules were found as main markers. Terpenoids (nine) and amino acids (eight) were the predominant compounds in this cluster. Triterpenoids of the classes oleae and dammarane were prevalent, i.e., celosin E and albasapogenin previously isolated from *Celosia argentea* and *Amaranthus hypochondriacus* (Amaranthaceae both), respectively (Wu et al., 2011) and ginsenoside isolated from *Panax ginseng* (Dou et al., 2001). Among the amino acids, peptides like Ac-D-Phe-βAla-L-His-D-Pro-NH2 (Edmundson et al., 1993), pacifibactin (Hardy & Butler, 2019), and aspartyl-tryptophan were identified. In the same cluster, an interesting marker found was a zearalenone derivative (3-(14,16-dihydroxy-3-methyl-1,7-dioxo-3,4,5,6,7,8,9,10-octahydro-1H-2-benzoxacyclotetradecin-15-yl)-N-[2-(1H-imidazol-1-yl)ethyl]-3-(3,4,5-trimethoxyphenyl) propenamide). To further characterize chemotype-specific biomarkers, a series of heatmaps were computed for each cluster and compiled in **supp info 6-8**. According to our findings, metabotyping is valuable in the classification of pseudocereals chemotypes, correlating the phylotranscriptomic clades of the family Amaranthaceae s.l.

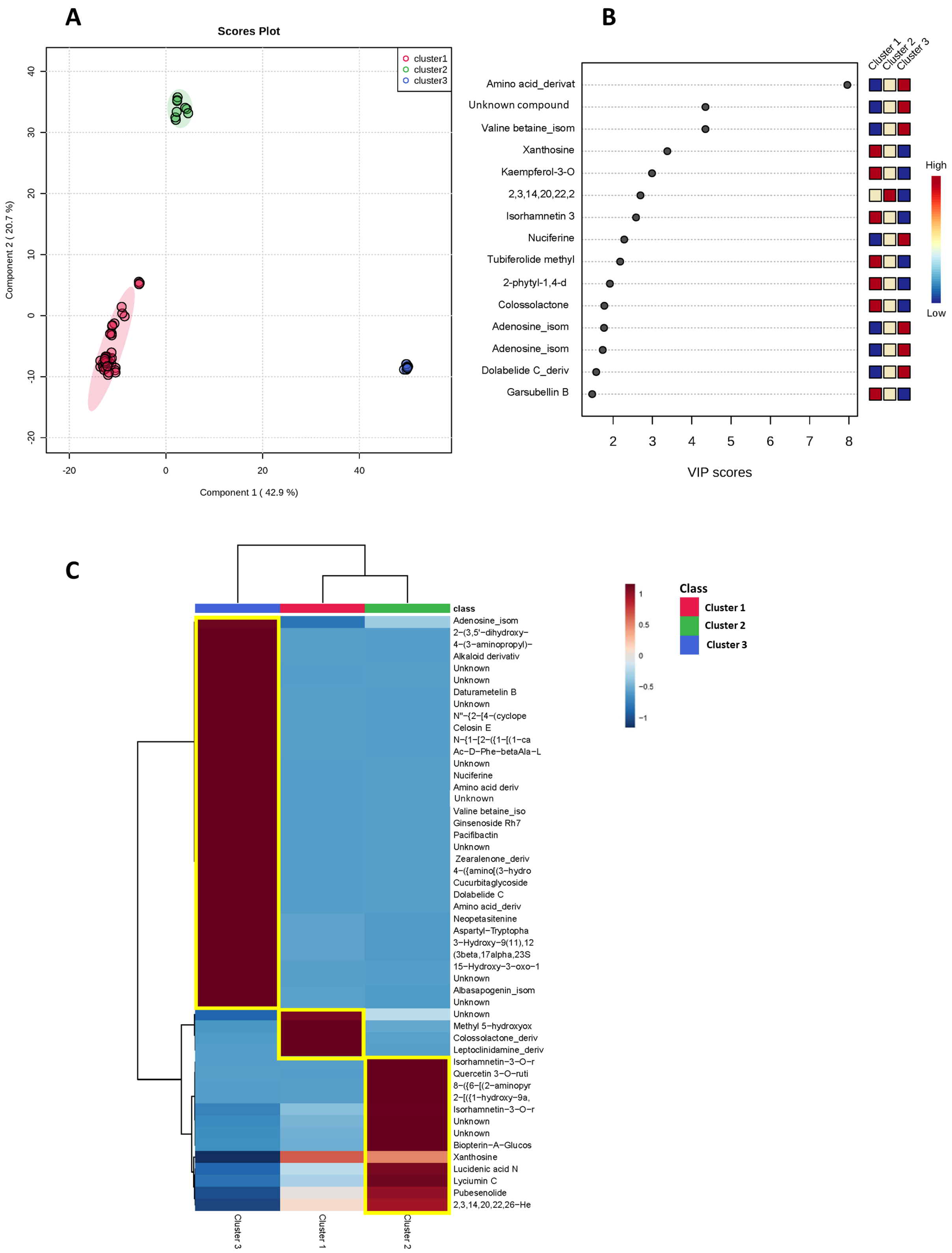


(**A**) Supervised PLSDA analysis clustering of the pseudocereal grains. (**B**) Most discriminating metabolites between clusters according to PLSDA based on the VIP values top 15. (**C**) Hierarchical clustering with the heatmap generated from the top 50 metabolites present in all varieties of pseudocereals. Clusters were grouped based on the HCA analysis showed in **Fig. 1-C**. Compounds were annotated using MSCleanR workflow.

5. Markers by cluster

| ID | *m/z* $[M+H]^+$ | RT | Formula | Compound annotated | Level of annotation | NPClassyfire (major classes) |
|---|---|---|---|---|---|---|
| | | | | **Cluster 1** | | |
| **138** | 337.573 | 5.079 | | Unknown compound | | |
| **60** | 222.076 | 4.357 | $C_{11}H_{11}NO_4$ | Methyl 5-hydroxyoxindole-3-acetate | generic | Alkaloid |
| **76** | 483.347 | 7.205 | $C_{31}H_{46}O_4$ | Colossolactone III_derivative | generic | Terpenoid |
| **146** | 362.146 | 6.128 | $C_{16}H_{19}N_5O_5$ | Leptoclinidamine B_derivative | | Amino acids and Peptides |
| | | | | **Cluster 2** | | |
| **6** | 625.176 | 6.011 | $C_{28}H_{32}O_{16}$ | isorhamnetin-3-O-rutinoside | family | Shikimates and Phenylpropanoids |
| **4** | 633.142[1] | 5.571 | $C_{27}H_{30}O_{16}$ | Quercetin 3-O-rutinoside | genus | Shikimates and Phenylpropanoids |
| **207** | 565.213 | 5.774 | $C_{30}H_{32}N_2O_9$ | 8-({6-[(2-aminopyridin-4-yl)methyl]-4,5,9-trihydroxy-1-(hydroxymethyl)-2-oxabicyclo[3.3.1]nonan-3-yl}oxy)-1-hydroxy-3-methyl-4a,9,9a,10-tetrahydroanthracene-9,10-dione | generic | Polyketides |
| **18** | 590.425 | 4.286 | $C_{37}H_{55}N_3O_3$ | 2-[({1-hydroxy-9a,11a-dimethyl-1H,2H,3H,3aH,3bH,4H,5H,7H,8H,9H,9aH,9bH,10H,11H,11aH-cyclopenta[a]phenanthren-7-ylidene}amino)oxy]-N-[2-(1,2,5-trimethyl-4-phenylpiperidin-4-yl)ethyl]acetamide | generic | Terpenoid |
| **41** | 625.176 | 5.449 | $C_{28}H_{32}O_{16}$ | isorhamnetin-3-O-rutinoside | | Shikimates and Phenylpropanoids |
| **178** | 460.720 | 7.26 | | Unknown compound | | |
| **29** | 781.151 | 5.223 | | Unknown compound | | |
| **152** | 400.146 | 2.885 | $C_{15}H_{21}N_5O_8$ | Biopterin-A-Glucoside | generic | Carbohydrate |
| **19** | 285.083 | 2.914 | $C_{10}H_{12}N_4O_6$ | Xanthosine | generic/in-house DB | Carbohydrate |
| **171** | 443.279[2] | 5.118 | $C_{27}H_{40}O_6$ | Lucidenic acid N_derivative | generic | Triterpenoi |
| **73** | 964.422 | 5.311 | $C_{49}H_{57}N_9O_{12}$ | Lyciumin C | generic | Amino acids and Peptides |
| **125** | 441.2[2] | 6.181 | $C_{28}H_{42}O_5$ | Pubesenolide | generic | Terpenoid |
| **9** | 445.295[2] | 5.781 | $C_{27}H_{44}O_7$ | 2,3,14,20,22,26-Hexahydroxycholest-7-en-6-one-form | family | Terpenoid |
| | | | | **Cluster 3** | | |
| **103** | 268.104 | 2.58 | $C_{10}H_{13}N_5O_4$ | Adenosine_isomer1 | family | Carbohydrate |
| **25** | 887.615 | 5.922 | $C_{54}H_{82}N_2O_8$ | 2-(3,5'-dihydroxy-4'-{6-[5-hydroxy-3-(2-methoxyethyl)-3,6-dimethyl-4-azabicyclo[9.3.1]pentadeca-1(14),6,11(15),12-tetraen-9-yl]-1-[(1-hydroxyheptan-2-yl)oxy]hepta-2,4,6-trien-2-yl}-8-(2-hydroxyethyl)-3-[2-(methylamino)ethyl]spiro[bicyclo[3.3.1]n | generic | Polyketides |

| | | | | | |
|---|---|---|---|---|---|
| | | | onane-2,1'-cyclopentan]-6-en-9-ylidene)propanal | | |
| **50** | 804.572 | 6.265 | $C_{52}H_{73}N_3O_4$ | 4-(3-aminopropyl)-7'-(cyclohexylmethyl)-26'-(2,5-dimethylhexyl)-18'-methyl-3H-3'-oxa-8',14'-diazaspiro[2-benzofuran-1,25'-heptacyclo,nonacosane]-4',15',21'(27')-triene-2',3-dione | generic | Alkaloid |
| **140** | 343.307 | 3.76 | $C_{18}H_{38}N_4O_2$ | alkaloid derivative | generic | Alkaloid |
| **22** | 964.419[3] | 7.452 | | Unknown compound | | |
| **75** | 1006.430 | 8.309 | | Unknown compound | | |
| **35** | 617.332 | 7.136 | $C_{34}H_{48}O_{10}$ | daturametelin B | generic | Terpenoid |
| **66** | 874.373[3] | 6.349 | | Unknown compound | | |
| **212** | 593.296 | 3.831 | $C_{32}H_{40}N_4O_7$ | N''-{2-[4-(cyclopentyloxy)-9-hydroxy-6-[4-(hydroxyamino)phenyl]-2-(2-hydroxypropan-2-yl)-5-oxo-2H,3H,5H,6H,7H,8H,9H-furo[3,2-b]xanthen-9-yl]ethyl}guanidine | generic | Other |
| **3** | 661.359[2] | 7.685 | $C_{36}H_{54}O_{12}$ | Celosin E | family | Terpenoid |
| **5** | 706.335 | 6.062 | $C_{39}H_{43}N_7O_6$ | N-{1-[2-({1-[(1-carbamoyl-2-phenylethyl)carbamoyl]-2-(1H-indol-3-yl)ethyl}carbamoyl)pyrrolidin-1-yl]-1-oxo-3-phenylpropan-2-yl}-5-oxopyrrolidine-2-carboxamide | generic | Amino acids and Peptides |
| **201** | 512.265 | 3.932 | $C_{25}H_{33}N_7O_5$ | Ac-D-Phe-betaAla-L-His-D-Pro-NH2 | generic | Amino acids and Peptides |
| **37** | 764.672 | 4.375 | | Unknown compound | | |
| **123** | 296.161 | 4.502 | $C_{19}H_{21}NO_2$ | Nuciferine | generic | Alkaloid |
| **48** | 174.149 | 2.687 | $C_9H_{19}NO_2$ | Amino acid derivative | generic | Amino acids and Peptides |
| **218** | 160.133 | 1.644 | | Unknown compound | | |
| **30** | 160.133 | 1.609 | $C_8H_{17}NO_2$ | Valine betaine | generic/intern DB | Amino acids and Peptides |
| **24** | 637.433 | 6.218 | $C_{36}H_{60}O_9$ | Ginsenoside Rh7 | generic | Terpenoid |
| **33** | 937.419 | 8.031 | $C_{35}H_{60}N_{12}O_{18}$ | Pacifibactin | generic | Amino acids and Peptides |
| **49** | 802.700 | 4.543 | | Unknown compound | | |
| **15** | 650.303 | 5.048 | $C_{35}H_{43}N_3O_9$ | Zearalenone derivative | generic | Other |
| **40** | 768.355 | 4.961 | $C_{35}H_{58}N_5O_{14}$ | 4-({amino[(3-hydroxypropyl)amino]methylidene}amino)-5-carboxylato-3-(2-{5-[(cyclopentyloxy)carbonyl]-3-ethenyl-2-{[4,4,5-trihydroxy-6-(hydroxymethyl)-3-[(methylazaniumyl)methoxy]oxan-2-yl]oxy}-3,4-dihydro-2H-pyran-4-yl}ethenyl)-1-(2-hydroxyethyl)piperidin-1-ium-3-yl | generic | Other |
| **69** | 926.432[2] | 6.926 | $C_{46}H_{65}N_5O_{16}$ | Cucurbitaglycoside A | generic | Terpenoid |
| **2** | 797.507 | 5.217 | $C_{43}H_{72}O_{13}$ | Dolabelide C_derivative | generic | Terpenoids |
| **23** | 733.335 | 5.279 | $C_{36}H_{44}N_8O_9$ | Amino acid derivative | generic | Amino acids and Peptides |
| **158** | 424.194 | 7.839 | $C_{21}H_{29}NO_8$ | Neopetasitenine | generic | Alkaloid |
| **134** | 320.125 | 4.168 | $C_{15}H_{17}N_3O_5$ | Aspartyl-Tryptophan | generic | Amino acids and Peptides |

| | | | | | | |
|---|---|---|---|---|---|---|
| **46** | 455.352 | 8.385 | $C_{30}H_{46}O_3$ | 3-Hydroxy-9(11),12-oleanadien-28-oic acid\|3-Hydroxy-9(11),12-oleanadien-28-oic acid_ 3-form\|3-Hydroxy-9(11),12-oleanadien-28-oic acid_ 3-form | family | Terpenoid |
| **68** | 471.311 | 7.375 | $C_{29}H_{42}O_5$ | (3beta,17alpha,23S)-17,23-Epoxy-3,29-dihydroxy-27-norlanosta-7,9(11)-diene-15,24-dione | generic | Terpenoid |
| **51** | 469.331 | 7.877 | $C_{30}H_{44}O_4$ | 15-Hydroxy-3-oxo-1,12-oleanadien-28-oic acid\|15-Hydroxy-3-oxo-1,12-oleanadien-28-oic acid_ 15-form | family | Terpenoid |
| **203** | 580.264[4] | 7.036 | | Unknown compound | | |
| **45** | 471.347 | 7.544 | $C_{30}H_{45}O_4$ | Albasapogenin | genus | Terpenoid |
| **200** | 510.727 | 7.914 | | Unknown compound | | |

[1] = $[M+Na]^+$; [2] = $[M+H-H_2O]^+$, [3] = $[2M+Na]^+$, [4] = $[M+2H]^{2+}$, DB = database

6. Heat map to cluster quinoa (cluster 1)

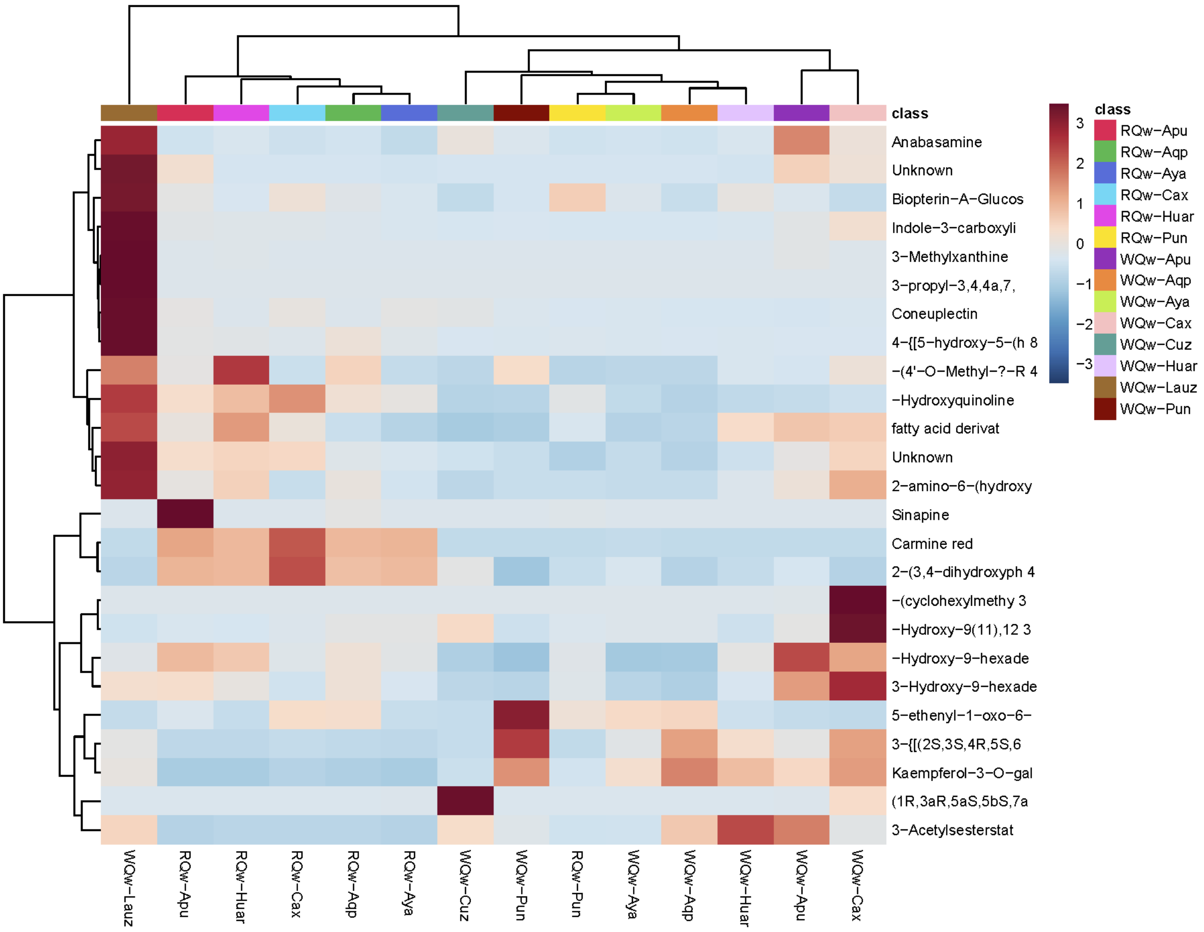

7. Heat map to cluster canihua (cluster 2)

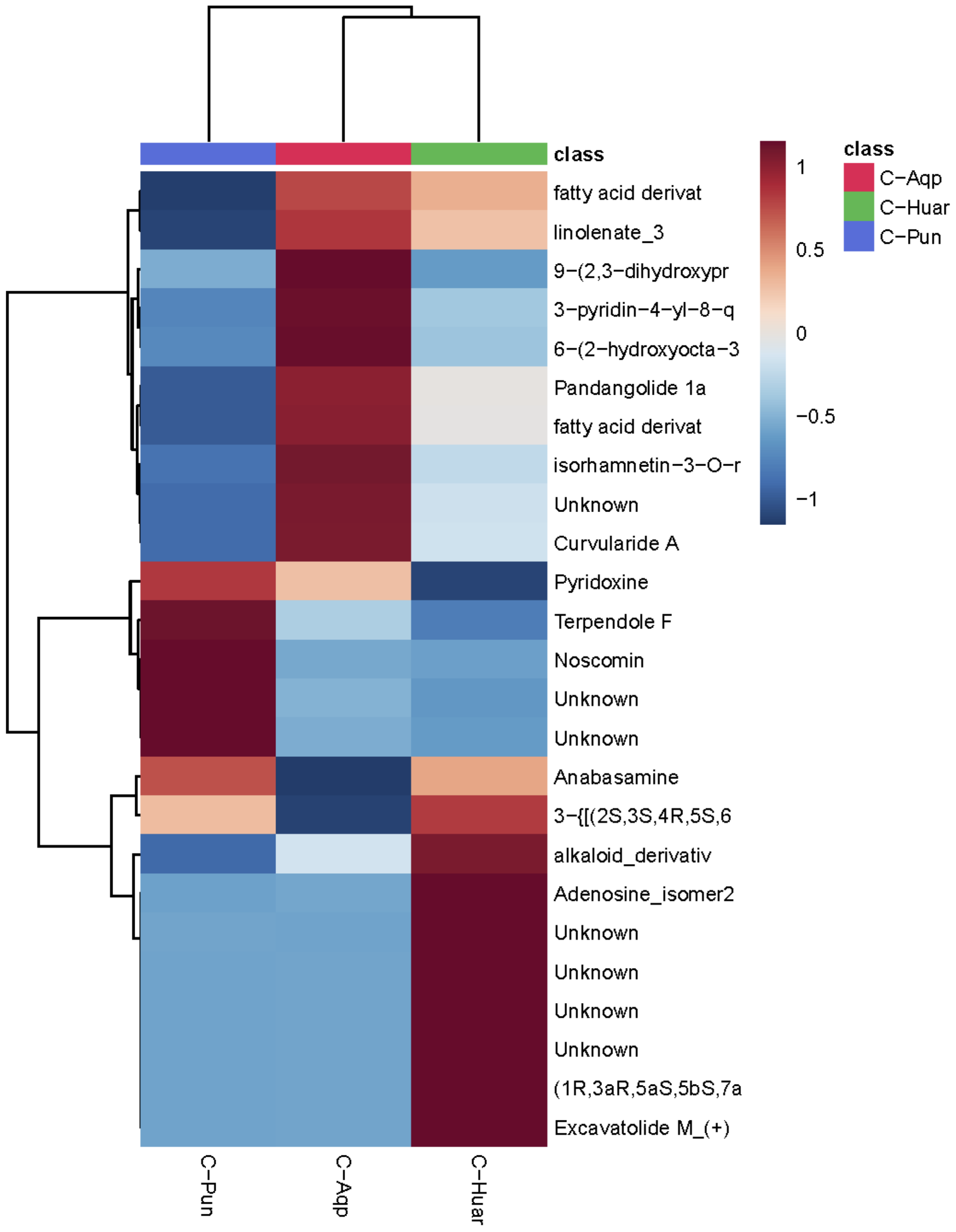

8. Heat map to cluster kiwicha (cluster 3)

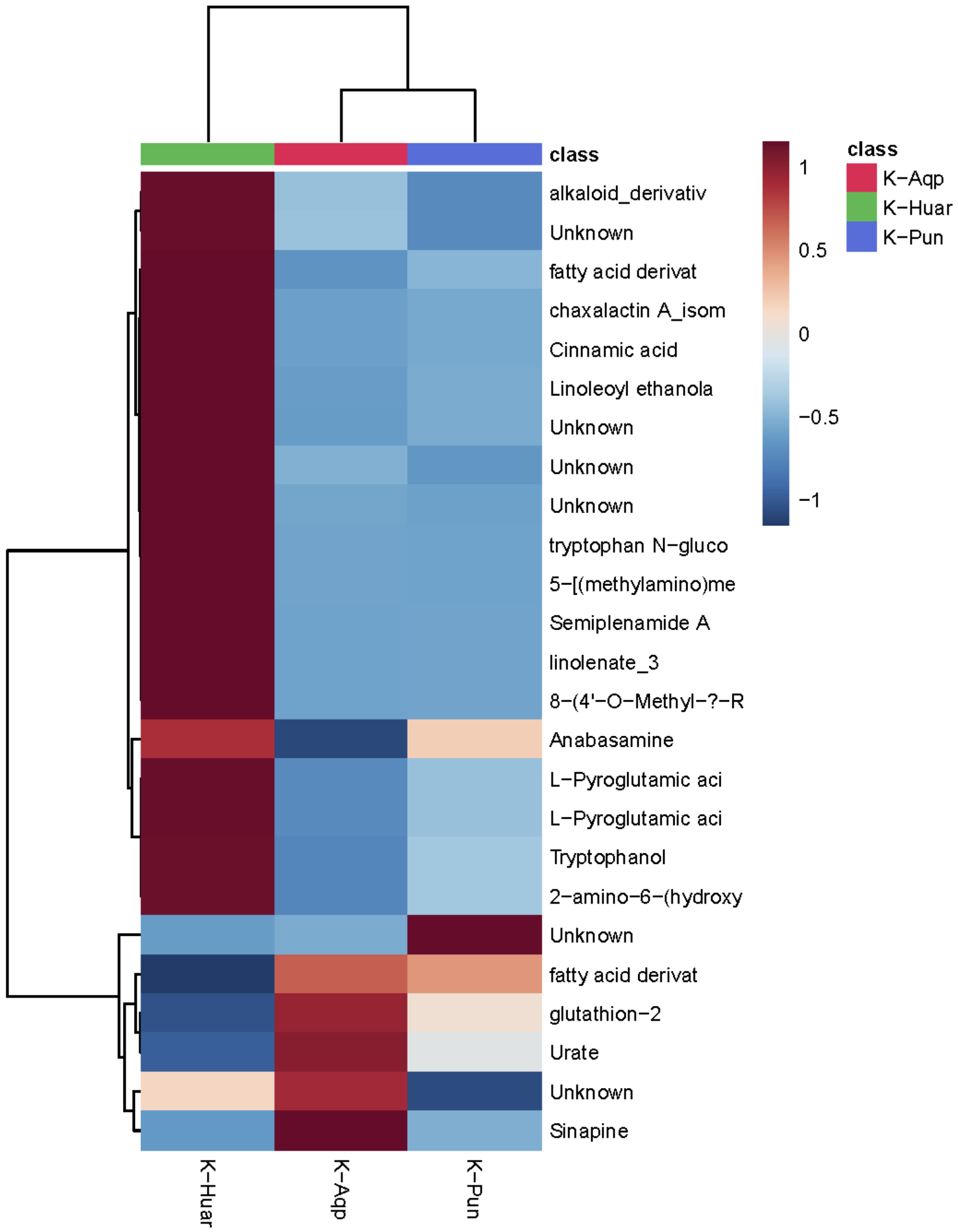

9. MS-MS pathway fragmentations for BEA and FumB1 by MRM

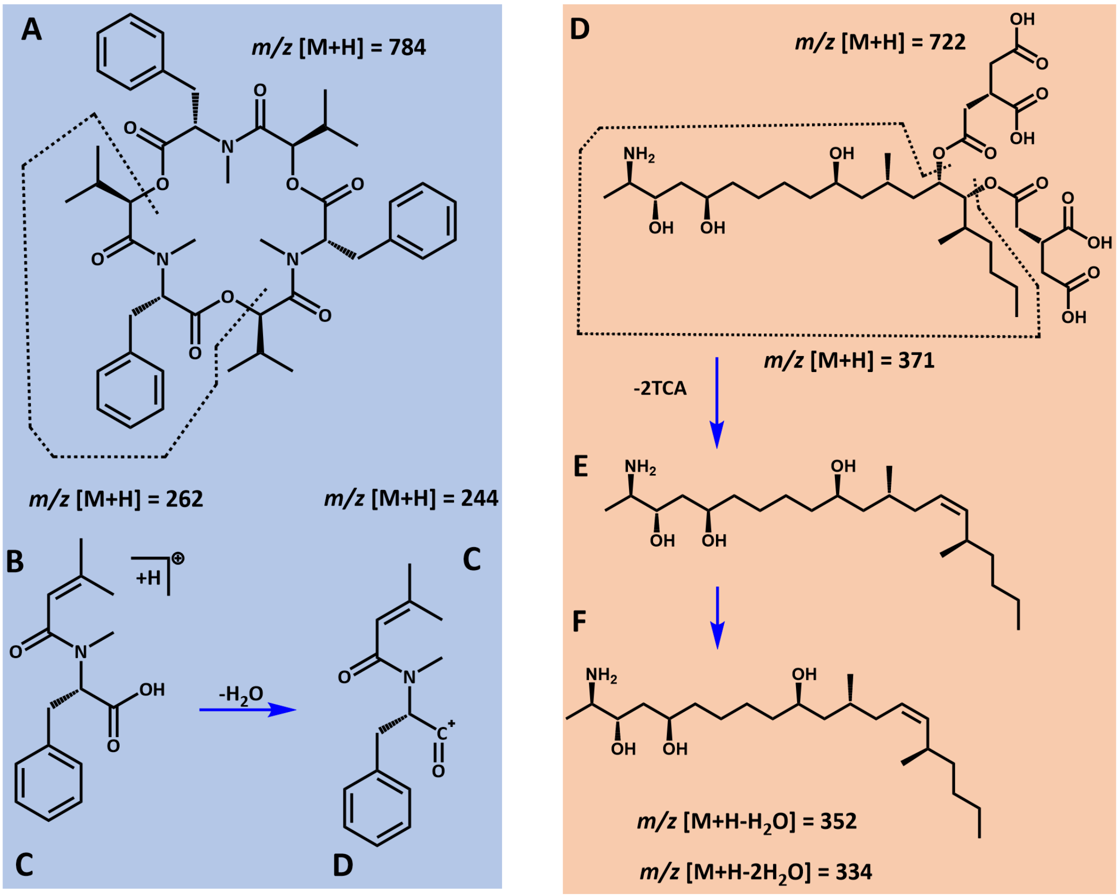


(**A**) Structure of BEA at *m/z* $[M+H]^+$ = 784. (**B**) Fragment of BEA containing phenylalanine residues at *m/z* $[M+H]^+$ = 262. (C) Fragment of (B) which then lost H2O to give *m/z* $[M+H]^+$ = 244 (fragment quantify by MRM). (**D**) Structure of FumB1 at *m/z* $[M+H]^+$ = 722. (**E**) Fragment corresponding to a loss of tricarballylic acid groups (TCA) from FumB1 at *m/z* $[M+H]^+$ = 371. (**F**) Loss of one or two molecules of water at *m/z* $[M+H-H2O]^+$ = 352 and *m/z* $[M+H-2H2O]^+$ = 334, respectively (fragments quantify by MRM).

10. Mycotoxins quantification from pseudocereal grains by MRM

| | **Beauvericin** | **$FumB_1$** |
|---|---|---|
| **Calibration curve equation** | 674890.7982x - 1596.9420 | 15452.7331x - 494.7162 |
| **Correlation coefficient value ($R^2$)** | 1.0 | 1.0 |
| **LOD (ng/mL)** | 0.005 | 0.05 |
| **LOQ (ng/mL)** | 0.019 | 0.19 |
| **Sample quantification, ng/mL in injected solution, mean ± SD (ng/g of powder ± SD)** | | |
| **WQun-Cuz1** | ND | ND |
| **WQun-Cuz2** | ND | ND |
| **WQw-Cuz** | ND | ND |
| **WQw-Aya** | ND | ND |
| **RQw-Aya** | ND | ND |
| **WQw-Apu** | ND | ND |
| **RQw-Apu** | ND | ND |
| **WQun-Cax** | 0.17 ± 0.01 (0.60 ± 0.04) | ND |
| **WQw-Cax** | 2.24 ± 0.24 (7.90 ± 0.85) | ND |
| **RQun-Cax** | < LOQ | ND |
| **RQw-Cax** | 0.13 ± 0.03 (0.45 ± 0.10) | ND |
| **WQun-Pun** | 0.019 ± 0.003 (0.05 ± 0.01) | ND |
| **WQw-Pun** | 0.04 ± 0.005 (0.11 ± 0.01) | ND |
| **RQun-Pun** | 0.04 ± 0.003 (0.15 ± 0.01) | ND |
| **RQw-Pun** | 0.02 ± 0.004 (0.07 ± 0.01) | ND |
| **K-Pun** | 0.03 ± 0.02 (0.12 ± 0.08) | ND |
| **C-Pun** | 0.39 ± 0.05 (1.71 ± 0.22) | ND |
| **WQun-Aqp** | 0.02 ± 0.003 (0.06 ± 0.01) | ND |
| **WQw-Aqp** | < LOQ | ND |
| **RQun-Aqp** | 0.03 ± 0.006 (0.10 ±0.02) | ND |
| **RQw-Aqp** | 0.3 ± 0.09 (1.06 ± 0.32) | < LOQ |
| **K-Aqp** | 0.06 ± 0.01 (0.22 ± 0.04) | < LOQ |
| **C-Aqp** | 0.03 ± 0.01 (0.11 ± 0.04) | < LOQ |
| **WQw-Huar** | < LOQ | < LOQ |
| **RQw-Huar** | 0.16 ± 0.02 (0.59 ± 0.07) | < LOQ |
| **K-Huar** | 0.03 ± 0.01 (0.13 ± 0.04) | ND |
| **C-Huar** | 0.03 ± 0.005 (0.14 ± 0.02) | < LOQ |
| **WQw-Lauz** | 0.44 ± 0.02 (1.26 ± 0.06) | < LOQ |
| **Wc-Bord** | 7.96 ± 0.41 (10.08 ± 0.52) | 0.47 ± 0.08 (0.60 ± 0.10) |

Wc-Bord = wheat contaminated from Bordeaux

11. Fungal strains isolated from Andean pseudocereal grains

| **Sample ID** | ***Species*** | **Family** | **Order** | | **Classe** | **ITS sequence** |
|---|---|---|---|---|---|---|
| WQun-Aqp_A | *Cladosporium complex cladosporioides* | Cladosporiaceae | Cladosporiales | Dothideomycetes | Ascomycetes | AMA18-A_ITS5 |
| WQun-Aqp_B | *Cladosporium complex cladosporioides* | Cladosporiaceae | Cladosporiales | Dothideomycetes | Ascomycetes | AMA18-B_ITS5 |
| WQun-Aqp_C | *Cladosporium complex cladosporioides* | Cladosporiaceae | Cladosporiales | Dothideomycetes | Ascomycetes | AMA18-C_ITS5 |
| WQun-Aqp_D | *Cladosporium complex cladosporioides* | Cladosporiaceae | Cladosporiales | Dothideomycetes | Ascomycetes | AMA18-D_ITS |
| WQun-Aqp_E | *Cladosporium complex cladosporioides* | Cladosporiaceae | Cladosporiales | Dothideomycetes | Ascomycetes | AMA18-E1_ITS |
| WQun-Cax_B | *Cladosporium complex cladosporioides* | Cladosporiaceae | Cladosporiales | Dothideomycetes | Ascomycetes | AMA08-B_ITS5 |
| WQw-Pun_A | *Cladosporium complex cladosporioides* | Cladosporiaceae | Cladosporiales | Dothideomycetes | Ascomycetes | AMA19-A_ITS5 |
| WQw-Aqp_A | *Cladosporium fusiforme* | Cladosporiaceae | Cladosporiales | Dothideomycetes | Ascomycetes | AMA08-A_ITS5 |
| WQun-Cax_A | *Didymella sp.* | Didymellaceae | Pleosporales | Dothideomycetes | Ascomycetes | AMA08-C_ITS |
| WQun-Cax_C | *Didymella sp.* | Didymellaceae | Pleosporales | Dothideomycetes | Ascomycetes | AMA08-D1_ITS |
| WQun-Cax_D | *Didymella sp.* | Didymellaceae | Pleosporales | Dothideomycetes | Ascomycetes | AMA01-B_ITS |
| WQun-Cuz1_B | *Epicoccum nigrum* | Didymellaceae | Pleosporales | Dothideomycetes | Ascomycetes | AMA02-L_ITS5 |
| WQun-Cuz2_L | *Epicoccum nigrum* | Didymellaceae | Pleosporales | Dothideomycetes | Ascomycetes | AMA02-A_ITS5 |
| RQun-Cax_D | *Alternaria cf. alternata* | Pleosporaceae | Pleosporales | Dothideomycetes | Ascomycetes | AMA02-B_ITS5 |
| WQun-Cuz2_A | *Alternaria cf. alternata* | Pleosporaceae | Pleosporales | Dothideomycetes | Ascomycetes | AMA02-D_ITS5 |
| WQun-Cuz2_B | *Alternaria cf. alternata* | Pleosporaceae | Pleosporales | Dothideomycetes | Ascomycetes | AMA03-A_ITS5 |
| WQun-Cuz2_D | *Alternaria cf. alternata* | Pleosporaceae | Pleosporales | Dothideomycetes | Ascomycetes | AMA03-G_ITS5 |
| WQw-Cuz_A | *Alternaria porri/solani** | Pleosporaceae | Pleosporales | Dothideomycetes | Ascomycetes | AMA02-C_ITS5 |
| WQw-Cuz_G | *Alternaria porri/solani** | Pleosporaceae | Pleosporales | Dothideomycetes | Ascomycetes | AMA02-F2_ITS5 |
| WQun-Cuz2_C | *Bipolaris victoriae* | Pleosporaceae | Pleosporales | Dothideomycetes | Ascomycetes | AMA17-C_ITS |
| WQun-Cuz2_F | *Bipolaris victoriae* | Pleosporaceae | Pleosporales | Dothideomycetes | Ascomycetes | AMA17-E_ITS5 |
| C-Pun_C | *Aspergillus amstelodami / montevidensis** | Aspergillaceae | Eurotiales | Eurotiomycetes | Ascomycetes | AMA17-F_ITS |
| C-Pun_E | *Aspergillus amstelodami / montevidensis** | Aspergillaceae | Eurotiales | Eurotiomycetes | Ascomycetes | AMA28-A_ITS |
| C-Pun_F | *Aspergillus amstelodami / montevidensis** | Aspergillaceae | Eurotiales | Eurotiomycetes | Ascomycetes | AMA28-B1_ITS |
| WQw-Lauz_A | *Aspergillus amstelodami / montevidensis** | Aspergillaceae | Eurotiales | Eurotiomycetes | Ascomycetes | AMA28-C_ITS |
| WQw-Lauz_B | *Aspergillus amstelodami / montevidensis** | Aspergillaceae | Eurotiales | Eurotiomycetes | Ascomycetes | AMA28-D1_ITS |
| WQw-Lauz_C | *Aspergillus amstelodami / montevidensis** | Aspergillaceae | Eurotiales | Eurotiomycetes | Ascomycetes | AMA28-G_ITS |

| | | | | | | |
|---|---|---|---|---|---|---|
| WQw-Lauz_D | *Aspergillus amstelodami / montevidensis** | Aspergillaceae | Eurotiales | Eurotiomycetes | Ascomycetes | AMA18-E2_ITS |
| WQw-Lauz_G | *Aspergillus amstelodami / montevidensis** | Aspergillaceae | Eurotiales | Eurotiomycetes | Ascomycetes | AMA18-F_ITS |
| WQun-Aqp_F | *Aspergillus amstelodami / montevidensis** | Aspergillaceae | Eurotiales | Eurotiomycetes | Ascomycetes | AMA07-A_ITS |
| WQun-Aqp_G | *Aspergillus amstelodami / montevidensis** | Aspergillaceae | Eurotiales | Eurotiomycetes | Ascomycetes | AMA21-A_ITS5 |
| RQw-Apu_A | *Aspergillus amstelodami / montevidensis** | Aspergillaceae | Eurotiales | Eurotiomycetes | Ascomycetes | AMA16-A_ITS5 |
| RQw-Aqp_A | *Aspergillus flavus* | Aspergillaceae | Eurotiales | Eurotiomycetes | Ascomycetes | AMA17-B_ITS5 |
| K-Pun | *Aspergillus fumigatus* | Aspergillaceae | Eurotiales | Eurotiomycetes | Ascomycetes | AMA17-D_ITS5 |
| C-Pun_B | *Aspergillus nidulans* | Aspergillaceae | Eurotiales | Eurotiomycetes | Ascomycetes | AMA14-A_ITS5 |
| C-Pun_D | *Aspergillus nidulans* | Aspergillaceae | Eurotiales | Eurotiomycetes | Ascomycetes | AMA14-C_ITS5 |
| RQun-Cax_C | *Aspergillus ochraceous* | Aspergillaceae | Eurotiales | Eurotiomycetes | Ascomycetes | AMA02-G_ITS |
| RQun-Pun_A | *Aspergillus sydowii* | Aspergillaceae | Eurotiales | Eurotiomycetes | Ascomycetes | AMA14-B_ITS5 |
| RQun-Pun_C | *Aspergillus welwitschiae* | Aspergillaceae | Eurotiales | Eurotiomycetes | Ascomycetes | AMA28-H_ITS |
| WQun-Cuz2_G | *Penicillium arizonense* | Aspergillaceae | Eurotiales | Eurotiomycetes | Ascomycetes | AMA26-A_ITS5 |
| RQun-Pun_B | *Penicillium chrysogenum* | Aspergillaceae | Eurotiales | Eurotiomycetes | Ascomycetes | AMA15-A_ITS |
| WQw-Lauz_H | *Penicillium cinnamopurpureum* | Aspergillaceae | Eurotiales | Eurotiomycetes | Ascomycetes | AMA15-B_ITS5 |
| K-Huar_A | *Penicillium crustosum* | Aspergillaceae | Eurotiales | Eurotiomycetes | Ascomycetes | AMA15-C_ITS5 |
| RQw-Pun_A | *Penicillium dipodomyis* | Aspergillaceae | Eurotiales | Eurotiomycetes | Ascomycetes | AMA15-D_ITS |
| RQw-Pun_B | *Penicillium dipodomyis* | Aspergillaceae | Eurotiales | Eurotiomycetes | Ascomycetes | AMA28-F_ITS5 |
| RQw-Pun_C | *Penicillium dipodomyis* | Aspergillaceae | Eurotiales | Eurotiomycetes | Ascomycetes | AMA03-B_ITS5 |
| RQw-Pun_D | *Penicillium dipodomyis* | Aspergillaceae | Eurotiales | Eurotiomycetes | Ascomycetes | AMA03-E_ITS5 |
| WQw-Lauz_F | *Penicillium sizovae* | Aspergillaceae | Eurotiales | Eurotiomycetes | Ascomycetes | AMA03-F_ITS5 |
| WQw-Cuz_B | *Talaromyces sp.* | Trichocomaceae | Eurotiales | Eurotiomycetes | Ascomycetes | AMA13_ITS5 |
| WQw-Cuz_E | *Talaromyces sp.* | Trichocomaceae | Eurotiales | Eurotiomycetes | Ascomycetes | AMA10-C_ITS5 |
| WQw-Cuz_F | *Talaromyces sp.* | Trichocomaceae | Eurotiales | Eurotiomycetes | Ascomycetes | AMA10-D_ITS5 |
| WQun-Cuz2_E | *Pyronema sp.* | Pyronemataceae | Pezizales | Pezizomycetes | Ascomycetes | AMA02-E_ITS5 |
| WQw-Aqp_B | *Eremothecium sinecaudum* | Saccharomycetaceae | Saccharomycetales | Saccharomycetes | Ascomycetes | AMA19-B_ITS5 |
| WQw-Cuz_C | *Trichoderma atroviride* | Hypocreaceae | Hypocreales | Sordariomycetes | Ascomycetes | AMA03-C_ITS5 |
| WQw-Cax_A | *Trichoderma longibrachiatum* | Hypocreaceae | Hypocreales | Sordariomycetes | Ascomycetes | AMA09-A_ITS5 |
| WQw-Cax_B | *Trichoderma longibrachiatum* | Hypocreaceae | Hypocreales | Sordariomycetes | Ascomycetes | AMA09-B_ITS5 |
| WQw-Cax_C | *Trichoderma longibrachiatum* | Hypocreaceae | Hypocreales | Sordariomycetes | Ascomycetes | AMA09-C_ITS5 |

| | | | | | | |
|---|---|---|---|---|---|---|
| WQun-Cuz2_K | *Purpureocillium lilacinum* | Ophiocordycipitaceae | Hypocreales | Sordariomycetes | Ascomycetes | AMA02-K_ITS5 |
| WQun-Cuz2_J | *Arthrinium sp.2* | Apiosporaceae | Xylariales | Sordariomycetes | Ascomycetes | AMA02-J_ITS5 |
| WQun-Cuz1_A | *Arthrinium sp.1* | Apiosporaceae | Xylariales | Sordariomycetes | Ascomycetes | AMA01-A_ITS5 |
| WQun-Cuz1_C | *Arthrinium sp.1* | Apiosporaceae | Xylariales | Sordariomycetes | Ascomycetes | AMA01-C_ITS5 |
| WQun-Cuz2_I | *Arthrinium sp.2* | Apiosporaceae | Xylariales | Sordariomycetes | Ascomycetes | AMA02-I_ITS5 |
| RQw-Pun_E | *Trametes versicolor* | Polyporaceae | Polyporales | Agaricomycetes | Basidiomycetes | AMA15-E_ITS5 |
| WQun-Cuz2_H | *Rhodotorula sp.* | Sporidiobolaceae | Sporidiobolales | Microbotryaomycetes | Basidiomycetes | AMA02-H_ITS |
| WQun-Cax_E | *Filobasidium magnum* | Filobasidiaceae | Filobasidilaes | Tremellomycetes | Basidiomycetes | AMA08-E_ITS |
| RQw-Huar | *Naganishia sp.* | Filobasidiaceae | Filobasidilaes | Tremellomycetes | Basidiomycetes | AMA25-B_ITS5 |
| WQun-Cuz1_D | *Ustilago sp.* | Ustilaginaceae | Ustilaginales | Ustilaginomycetes | Basidiomycetes | AMA01-D_ITS |
| C-Pun_A | *Lichtheimia (Absidia) corymbifera* | Lichtheimiaceae | Mucorales | Mucoromycetes | Mucoromycetes | AMA17-A_ITS5 |
| C-Aqp_A | *Rhizopus oryzae/delemar** | Rhizopodaceae | Mucorales | Mucoromycetes | Mucoromycetes | AMA23-A_ITS5 |
| K-Aqp_A | *Rhizopus oryzae/delemar** | Rhizopodaceae | Mucorales | Mucoromycetes | Mucoromycetes | AMA22-B_ITS5 |
| RQw-Cax_A | *Rhizopus oryzae/delemar** | Rhizopodaceae | Mucorales | Mucoromycetes | Mucoromycetes | AMA11-A_ITS5 |
| RQun-Aqp_A | *Rhizopus oryzae/delemar** | Rhizopodaceae | Mucorales | Mucoromycetes | Mucoromycetes | AMA20-A_ITS5 |
| RQun-Cax_A | *Rhizopus oryzae/delemar** | Rhizopodaceae | Mucorales | Mucoromycetes | Mucoromycetes | AMA10-A_ITS5 |
